# Cysteine Restriction-Specific Effects of Sulfur Amino Acid Restriction on Lipid Metabolism

**DOI:** 10.1101/2022.05.14.491847

**Authors:** Sailendra N Nichenametla, Dwight AL Mattocks, Diana Cooke, Vishal Midya, Virginia L Malloy, Wilfredo Mansilla, Bente Øvrebø, Cheryl Turner, Nasser E Bastani, Jitka Sokolová, Markéta Pavlíková, John P Richie, Anna K Shoveller, Helga Refsum, Thomas Olsen, Kathrine J Vinknes, Viktor Kožich, Gene P Ables

## Abstract

Decreasing dietary intake of methionine exerts robust anti-adiposity effects in rodents but modest effects in humans. Since cysteine can be synthesized from methionine, animal diets are formulated by decreasing methionine and eliminating cysteine. Such diets exert both methionine restriction (MR) and cysteine restriction (CR), i.e., sulfur amino acid restriction (SAAR). Contrarily, human diets used in clinical studies did not eliminate cysteine and thus might have exerted only MR. Epidemiological studies positively correlate body adiposity with plasma cysteine but not with methionine, suggesting that CR, but not MR, is responsible for the antiadiposity effects of SAAR in rodents. Whether this is true, and if so, the underlying mechanisms are unknown. Using multiple diets with variable concentrations of methionine and cysteine, we demonstrate that the anti-adiposity effects of SAAR are due to CR. CR increased serinogenesis (serine biosynthesis from non-glucose substrates) by diverting substrates from glyceroneogenesis, essential for fatty acid/triglyceride cycling. Molecular data suggest that the CR results in glutathione depletion, which induces Nrf2 and downstream targets Phgdh (the serine biosynthetic enzyme) and Pepck-M. Using multiple mouse models, we show that the magnitude of SAAR-induced changes in molecular markers depends on dietary fat concentration (60 fat>10% fat), gender (males>females), and age-at-onset (young>adult). Our findings are translationally relevant as we found a negative correlation of plasma serine with triglycerides and metabolic syndrome criteria in a cross-sectional epidemiological study. SAAR-like diets with high polyunsaturated fatty acids increased plasma serine in a short-term human feeding study. Serinogenesis might be a potential target to correct hypertriglyceridemia.

## Introduction

Dysregulated lipid metabolism is a common etiological factor in obesity, metabolic syndrome (MetS), and type 2 diabetes (Stout, Justice, Nicklas, & Kirkland, 2017). It also increases the risk of inflammatory diseases and cancers (Catalan, Gomez-Ambrosi, Rodriguez, & Fruhbeck, 2013; Gutierrez, Puglisi, & Hasty, 2009; Salvestrini, Sell, & Lorenzini, 2019). Improving lipid metabolic homeostasis would not only ameliorate these diseases but also extends overall healthspan. Preclinical studies report that lowering the dietary intake of the sulfur amino acid methionine (Met) in the absence of cysteine (Cys) (sulfur amino acid restriction - SAAR) improves lipid metabolism even with *ad libitum* access to the diet (X. Zhou et al., 2016). SAAR not only makes mice resistant to diet-induced obesity but also induces weight loss in obese mice within two weeks (Ables, Perrone, Orentreich, & Orentreich, 2012; Cooke et al., 2020). SAAR-induced benefits in lipid metabolism include decreases in body weight, fat mass, plasma triglycerides, and leptin; and increases in plasma adiponectin and fibroblast growth factor 21 (Fgf21). The high reproducibility and the strong effect of SAAR on rodent lipid metabolism spurred clinical studies to combat obesity and MetS (Olsen et al., 2020; Olsen et al., 2018; Plaisance et al., 2011). However, the few clinical studies conducted so far show that its impact on humans is very modest compared to the effects of SAAR in animals. Although the molecular signature of SAAR-induced changes in lipid metabolism, including altered expression of hormones (insulin-like growth factor 1, Fgf21, adiponectin, and leptin), enzymes (fatty acid synthase, acetyl-CoA carboxylase, Stearoyl-CoA desaturase, carnitine palmitoyltransferase1, and adipose triglyceride lipase), and metabolites (S-adenosylmethionine, S-adenosylhomocysteine, carnitine, ceramide, and sphingosine) is well-characterized in rodents, the biochemical events that drive these changes remain unknown (X. Zhou et al., 2016). Understanding the biochemistry of SAAR-induced changes in rodents is critical for its translation to humans.

Met and Cys are the two proteinogenic sulfur amino acids (SAA) with distinct biological roles. Although the metabolic requirement of Cys is essential, its dietary requirement is dispensable, as healthy adult animals and humans can convert Met to Cys through the transsulfuration pathway (Womack, Kemmerer, & Rose, 1937). Accordingly, the requirement of total SAA, i.e., Met and Cys together, in synthetic rodent diets is usually met by providing only Met. Most animal studies formulated SAAR diets by decreasing the Met content of the diet by approximately 80% (0.86% w/w dry matter content in control diets and 0.12% - 0.17% w/w dry matter content in the SAAR diet) and eliminating Cys. This approach is justified if the objective is to investigate the combined effect of low SAA on a response variable but not their discrete effects. Although healthy rodents have a well-functioning transsulfuration pathway, the Met concentration in the SAAR diet is too low to meet growth and metabolic demands for Cys (Reeves, Nielsen, & Fahey, 1993). This insufficiency implies that rodents on SAAR undergo both Met restriction (MR) and Cys restriction (CR). Whether the SAAR-induced changes are due to the combined or discrete effects of MR and CR remains unknown. Understanding the distinct roles of individual SAA is critical since multiple epidemiological studies and a limited number of laboratory studies suggest that the SAAR effects on lipid metabolism are due to CR but not MR.

Most laboratory studies use SAAR diets devoid of Cys. However, a few studies supplemented SAAR diets with Cys. In one study, male F344 rats were fed the typical SAAR diet (0.17% w/w Met without Cys) and the SAAR diet supplemented with 0.5% w/w Cys (Elshorbagy et al., 2011; Perrone et al., 2012). With Cys supplementation, SAAR-induced decrease in body weight, adipose depot weights, plasma triglycerides, leptin, total cholesterol, and LDL cholesterol were reversed. In another study, mature C3H/HeH mice were fed two diets, each with different concentrations of Cys (0.07% w/w and 0.87% w/w) but with the same concentration of Met (0.29% w/w). Mice on the diet with high Cys had greater body weights, adipose depot weights, hepatic triglycerides, plasma triglycerides, and LDL-cholesterol than those on low Cys (Elshorbagy, Church, et al., 2012). Large-scale epidemiological studies show that plasma Cys, but not Met, is positively associated with body mass index and fat mass independent of caloric intake and physical activity, thus rendering support to the preclinical studies (Elshorbagy, Kozich, Smith, & Refsum, 2012; Elshorbagy et al., 2008; Elshorbagy, Smith, Kozich, & Refsum, 2012). Despite these data, one cannot ascertain changes in lipid metabolic markers to CR alone as the reversal of the SAAR-induced changes upon Cys supplementation could also be due to the biological abundance of Met caused by the sparing effect of dietary Cys on the transsulfuration pathway.

In the current study, we demonstrate that MR and CR exert discrete effects on SAAR-induced changes by conducting depletion-repletion bioassays. First, a cohort of male Fischer 344 (F344) rats was fed four diets, all replete with 0.5% w/w Cys, but formulated with progressive MR. Based on data from the first cohort, we provided the second cohort of rats with five diets, all with the same low Met concentration (0.07% w/w) but with progressively restricted concentrations of Cys. SAAR-induced changes by MR- and CR-diets were compared with those induced by the typical SAAR (0.17% w/w Met) and control diets (CD, 0.86% w/w Met), both of which were used in our previous studies and were devoid of Cys (Nichenametla, Mattocks, & Malloy, 2020). To understand the biochemical and molecular mechanisms of CR-specific changes in adipose metabolism, we quantified transcriptional and translational changes in multiple genes and proteins. We also investigated if the CR-specific effects on adipose metabolism are dependent on gender, age-at-onset (AAO), and dietary fat content in C57BL/6J mice. Later, we tested the relevance of our preclinical findings to humans. We reanalyzed data from two short-term SAAR feeding studies. In addition, we analyzed biospecimens and data from a previously conducted population study in individuals with different degrees of MetS.

## Results

### 1. Methionine restriction and cysteine restriction exert discrete effects on morphometry and plasma hormone concentrations

Due to the conversion of Met to Cys through the transsulfuration pathway, effectuating MR without CR or *vice versa* in the context of the SAAR diet is not straightforward. We overcame this problem by taking a stepwise approach. First, to obviate the need for Met conversion to Cys, we fed a cohort of rats four different MR diets, all replete with Cys (0.5% w/w), but progressively restricted in Met (MR1 - 0.17%, MR2 - 0.10%, MR3 - 0.07%, and MR4 - 0.05% w/w). Two other groups of rats were fed the standard CD (0.86% w/w Met without Cys) and SAAR (0.17% w/w Met without Cys) diets. All diets had the same caloric density and macronutrient concentrations (Supplementary Tables 1a and 1b). SAAR-induced changes in all markers were in agreement with previous studies, i.e., compared to the rats on CD, those on SAAR had slower growth rates (Figure 1a, ratio of mean values of SAAR to CD [SAAR/CD] - 0.31; p <0.0001), lower Igf1 (Figure 1c, SAAR/CD - 0.60; p<0.0001), and leptin (Figure 1e, SAAR/CD - 0.44; p<0.0001), and higher food consumption (Figure 1b, SAAR/CD - 1.19; p <0.0001), Fgf21 (Figure 1d, SAAR/CD - 4.86; p<0.01), and adiponectin (Figure 1f, SAAR/CD - 3.23; p <0.0001) (Nichenametla et al., 2020).

**Figure 1.**
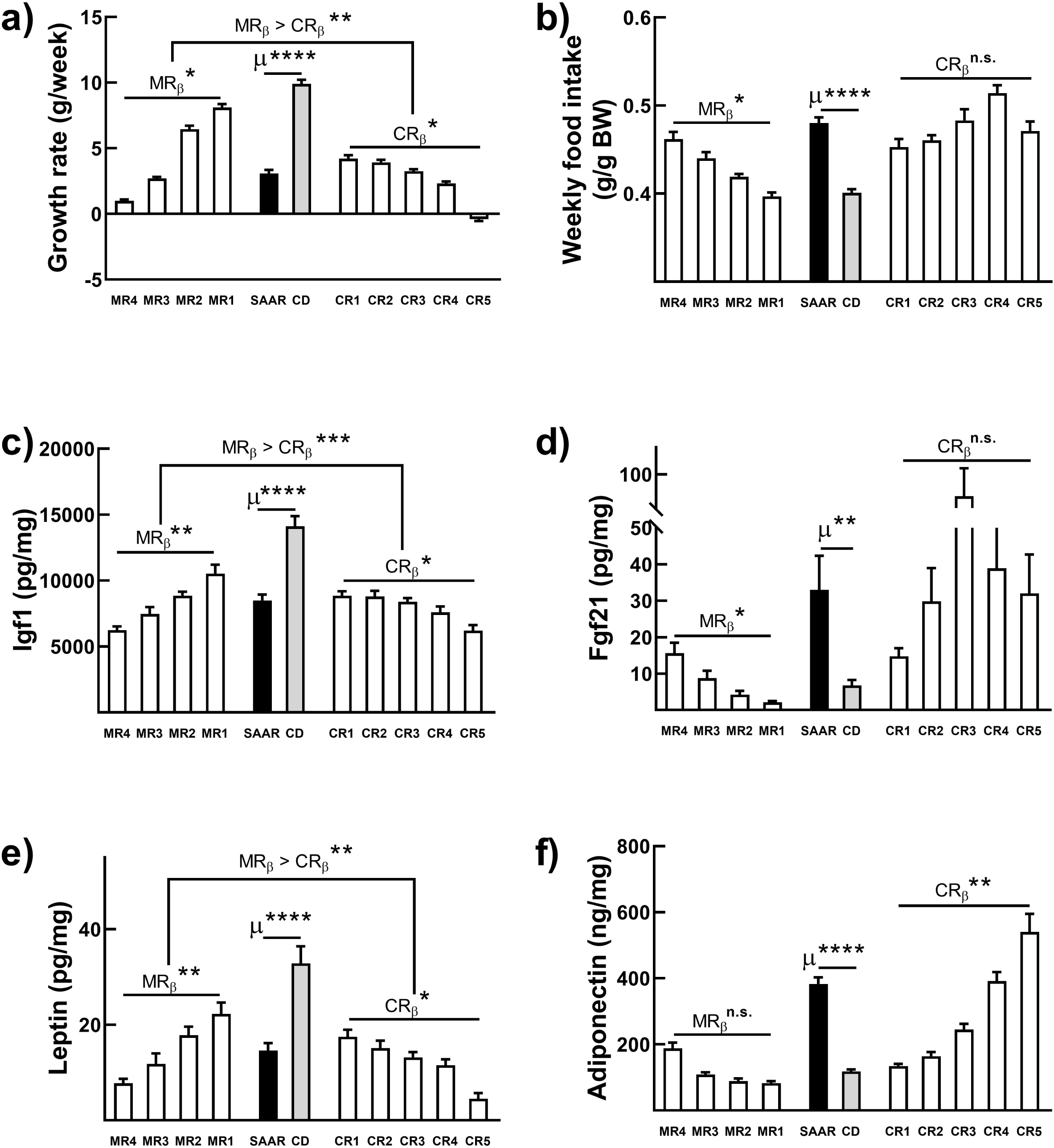

Whether any of the six SAAR-induced changes were dependent on MR or not was determined by calculating the regression coefficients (MR_β_) for each phenotype (response) with the increasing dose of MR (MR1, MR2, MR3, and MR4) Growth rates (MR_β_ [-0.89]; p < 0.05, Figure 1a), food intake (Figure 1b, MR_β_ [0.86]; p < 0.05), Igf1 (MR_β_ [-0.59]; p < 0.01, Figure 1c), Fgf21 (Figure 1d, MR_β_ [0.55]; p < 0.05), and leptin (MR_β_ [-0.50]; p < 0.01, Figure 1e) exhibited a strong dose-response to MR, while no dose-response was observed for adiponectin (Figure 1f). The dose of MR that achieved a similar effect size in measured endpoints as observed with the typical SAAR diet (0.17% Met without Cys) was considered the minimal obligatory dose of methionine (MOM). We expect that the Met flux to transsulfuration pathway would be minimal at this dose even upon restricting dietary Cys. Based on equivalence testing between the SAAR diet and each of the four MR diets, we considered MR3 (0.07% with 0.5% Cys) as the MOM, the dose at which Met was provided to examine the discrete effect of CR (Supplementary Figure 1). The second cohort of rats was fed five different diets, all consisting of Met at the MOM level, but each with progressively restricted Cys (CR1 - 0.5%, CR2 - 0.25%, CR3 - 0.12%, CR4 - 0.06%, and CR5 - 0.03% w/w). MR-dependent markers were either non-responsive to CR (food intake and Fgf21, Figures 1b and 1d) or less responsive (growth rate - CR_β_ [-0.34]; p < 0.05, Igf1 - CR_β_ [-0.19]; p < 0.05, and leptin CR_β_ [-0.25]; p <0.05, Figures 1a, 1c, and 1e) to CR than MR. Plasma adiponectin, however, which did not respond to MR, exhibited a strong dose-response to CR (Figure 1f, CR_β_ [0.62]; p < 0.01). Overall, our data confirm that among the changes induced by the SAAR diet, changes in growth rates, food intake, plasma Igf1, and leptin were MR-dependent (Figures 1a-1e). In contrast, plasma adiponectin was CR-dependent (Figure 1f). Among the five CR diets, CR4 yielded a similar effect size for adiponectin as for the SAAR diet (Supplementary Figure 2). For reasons we cannot explain, the dose-response of Fgf21 appears to be parabolic. Previous studies demonstrated that Fgf21 is extremely sensitive to fasting and varies up to 250-fold in humans (Galman et al., 2008). Considering the larger standard errors of the mean in rats on CR diets than those on MR diets, it would be interesting to investigate whether Cys contributes to this variation.

The MR1 diet in this study yielded similar results on the same endpoints as a previous study using an identical diet (0.17% Met; 0,5% Cys) (Elshorbagy et al., 2011). In that study, based on the ability of Cys to reverse the SAAR-induced phenotypes, the authors suggested that CR was responsible for their induction. Our study shows that with increasing degrees of MR, MR-dependent endpoints progressively recapitulated even when supplemented with 0.5% Cys, i.e., in MR2, MR3, and MR4 diets. This phenomenon indirectly establishes that the reversal of certain SAAR-induced changes observed in the previous report was a result of the sparing effect of dietary Cys on the transsulfuration pathway. Another report suggested that the ideal concentration range of Met in the SAAR diet was between 0.17% and 0.25% and that levels below 0.12% invoke amino acid deprivation responses (Forney, Wanders, Stone, Pierse, & Gettys, 2017). Although this interpretation is accurate, our findings show that when supplemented with 0.5% Cys, Met concentration in the SAAR diet can be lowered to 0.05% without any adverse effects. All rats were healthy and active throughout the study and showed no signs of amino acid deficiency (lethargy, hair loss, and decreased food intake).

### 2. Methionine and cysteine restrictions induce distinct changes in plasma amino acid profiles

We previously reported that SAAR decreases hepatic protein synthesis rates by 33%, but the metabolic fate of the amino acids not utilized for protein synthesis is unknown (Nichenametla, Mattocks, Malloy, & Pinto, 2018). After deamination/transamination, the carbon skeleton of amino acids can affect lipid metabolism through anaplerosis. We and others reported that individuals on a low Met diet had higher levels of plasma oxaloacetate, indicating changes in the TCA cycle (Gao et al., 2019; Martinez-Reyes & Chandel, 2020). We quantified plasma amino acids to determine if they might mediate some of the SAAR-induced changes in adipose metabolism and if MR and CR discretely affect their concentrations. Contrary to the strong association between plasma branched-chain amino acids and adiposity, SAAR did not alter plasma Leu, Ile, and Val, despite its anti-adiposity effects (data not presented) (Siddik & Shin, 2019). However, of the 20 proteinogenic amino acids identified in our analysis, SAAR increased the levels of Glu (ratio of the mean concentrations of SAAR to CD [SAAR/CD] -1.15, p<0.05), Lys (SAAR/CD-1.16, p<0.01), His (SAAR/CD-1.16, p<0.01), Ser (SAAR/CD-1.97, p<0.0001), and Thr (SAAR/CD-2.65, p<0.0001), and decreased the levels of Phe (SAAR/CD-0.81, p<0.0001) and Gly (SAAR/CD-0.82, p<0.01) (Figures 2a-2i). It is noteworthy that despite significant differences between CD and SAAR, Glu, Gly, and Lys were not dose-responsive to either MR or CR (Figures 2a-2c). On the other hand, Phe (CR_β_ [-0.51]; p < 0.01), His (CR_β_ [0.69]; p < 0.05), Ser (CR_β_ [0.55]; p < 0.05), Thr (CR_β_ [0.54]; p < 0.05), and Trp (CR_β_ [-0.62]; p < 0.01) were dose-responsive to CR but not MR (Figures 2d-2i). Despite there being no significant change between CD and SAAR, Met exhibited a dose response to CR (Figure 2e, CR_β_ [-0.41], p < 0.01). Since many studies reported that Ser alters lipid metabolism, particularly lipid profiles in pathological conditions, we focused our additional analyses on Ser (Esch et al., 2020; Gantner et al., 2019; Gao et al., 2018; Muthusamy et al., 2020).

**Figure 2.**
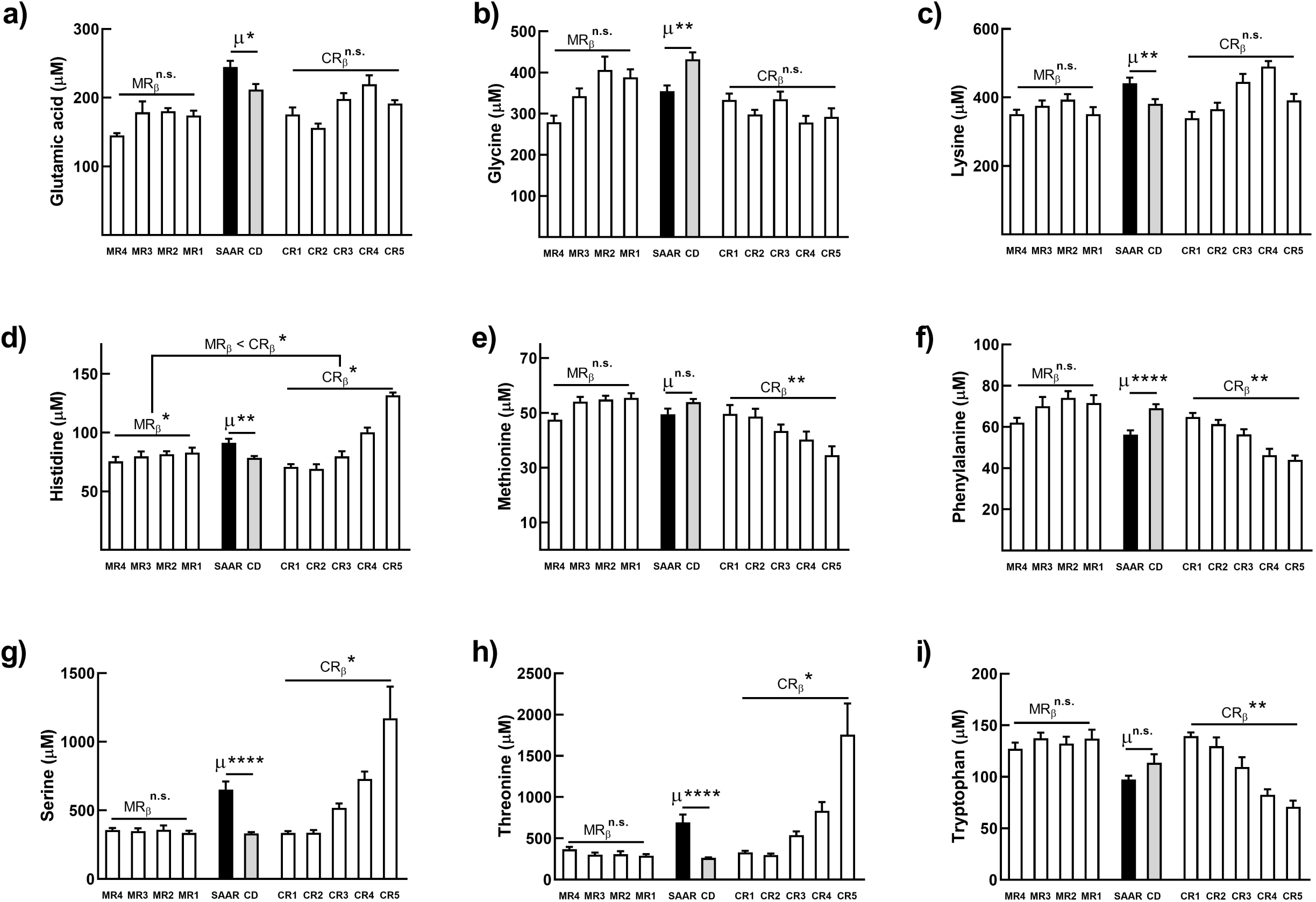

### 3. Cysteine restriction but not methionine restriction induces hepatic *de novo* serine biosynthesis

To understand if and how increased plasma Ser is involved in SAAR-induced changes in lipid metabolism, we wanted to identify the organs involved. Higher plasma Ser could be a consequence of increased biosynthesis, decreased utilization, or both. Multiple reports show that the kidney and liver are the major tissues that affect plasma Ser concentrations (Brosnan & Hall, 1989; Lowry, Hall, Hall, & Brosnan, 1987). Since we did not harvest kidneys, we quantified hepatic Ser. Compared to CD, SAAR increased hepatic Ser concentration about 2-fold (p<0.0001, Figure 3a). Our findings agree with a recent report that shows higher hepatic Ser in SAAR-fed mice (Stone et al., 2021). Similar to the CR-dependent increase in plasma Ser, the increase in hepatic Ser was also dependent on CR (CR_β_ [0.59]; p < 0.01) but not on MR. To confirm if the increase in hepatic Ser is due to its *de novo* biosynthesis, we probed for the transcript levels of the rate-limiting enzyme for Ser biosynthesis, phosphoglycerate dehydrogenase (Phgdh). Compared to CD, hepatic *Phgdh* expression was approximately 6- to 9-fold higher in SAAR (Figures 3b and 3c, p < 0.01). A robust dose-response was found in *Phgdh* levels with CR but not with MR (Figures 3b and 3c, CR_β_ [0.51]; p < 0.05). In agreement with our findings on *Phgdh*, previous studies based on nuclear run-off assays show that dietary protein regulates *Phgdh* mRNA post-transcriptionally. In particular, higher concentrations of Cys, but not of Met, destabilized *Phgdh* mRNA (Achouri, Robbi, & Van Schaftingen, 1999). To confirm whether increases in mRNA translate to protein levels, we probed for hepatic Phgdh in rats fed diets with the highest and lowest SAA levels (MR1, MR4, CR1, and CR5). MR1 and MR4 had very low but similar levels of Phgdh (Figure 3d). No bands were detected in CR1, but the band intensity in CR5 was greater than that in the SAAR (Figure 3d). Following this, to check if Phgdh exhibits a dose-response to CR, we probed for hepatic Phgdh in all CR diet groups. Phgdh was dose-responsive to CR (Figure 3e, CR_β_ [0.43]; p < 0.01). These data strongly suggest that SAAR increases the *de novo* hepatic Ser biosynthesis and that this effect is specifically due to CR. A previous study reported that hepatic Ser biosynthesis was not appreciable in rodents when fed diets with normal protein concentration but was amplified on protein-restricted diets (Kalhan et al., 2011). Considering that all the diets in our study had similar total amino acid content, our findings implicate that some of the benefits induced by protein-restricted diets could be due to low SAA.

**Figure 3.**
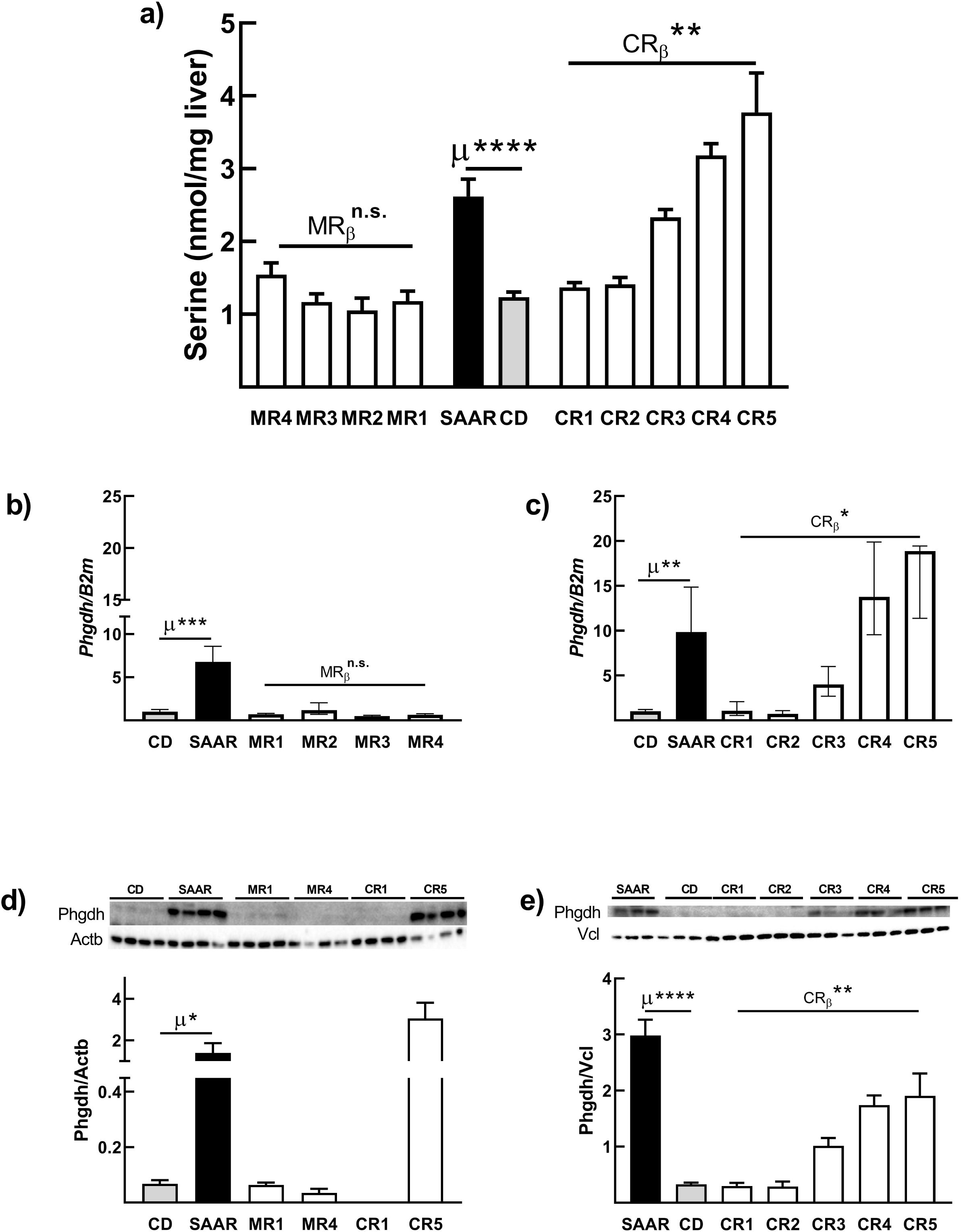

### 4. 3-Phosphoglycerate, the substrate for serine biosynthesis, is derived from mitochondrial oxaloacetate

To sustain the increase in hepatic *de novo* Ser biosynthesis, a constant supply of the substrate 3-phosphoglycerate (3PG) is required. The canonical pathway for Ser biosynthesis under normal conditions is the “phosphorylated pathway,” in which the substrate 3PG is derived from glucose through glycolysis (Murtas, Marcone, Sacchi, & Pollegioni, 2020). If CR-induced Ser biosynthesis is through glycolysis, animals on CR diets should exhibit a dose-dependent decrease in blood glucose. However, our data show that, unlike plasma Ser, fasting blood glucose responded to MR but not CR (Figure 4a, MRβ [-0.68]; p < 0.5), which indicates that glycolysis may not be contributing to Ser biosynthesis. Another likely reason for the non-glycolytic origin of Ser is that hepatic glycolysis is minimal during post-absorptive states, such as the prandial state in which plasma and tissues were obtained. Since 3PG can also be derived from gluconeogenesis, we probed for the mRNA expression of two enzymes that regulate this pathway, phosphoenolpyruvate carboxykinase-C (Pepck-C, coded by gene *Pck1*), the rate-limiting enzyme, and glucose-6-phosphatase-c (G6pc, the catalytic subunit of G6p), the terminal enzyme which releases glucose from glucose-6-phosphate. SAAR did not alter the mRNA expression of either of these enzymes (Figures 4b, 4c, 4d, and 4e), also suggesting that gluconeogenesis may not be responsible for CR-induced Ser biosynthesis. To our knowledge, there is no other major source of 3PG besides glycolysis/gluconeogenesis pathways. A recent study reported coordination between the mitochondrial form of Pepck, Pepck-M (coded by the gene *Pck2*), and Ser biosynthesis (Brown et al., 2016). To check if this coordination occurs in the liver in the context of SAAR, we probed for hepatic *Pck2* mRNA expression. We not only found that SAAR increased *Pck2* mRNA expression by at least 3.5-fold (Figures 4f and 4g, p <0.0001) but also that, similar to the CR-dependent increase in hepatic Ser and Phgdh, *Pck2* mRNA exhibited a strong dose-response to CR (CR_β_ < 0.05) but not to MR. In addition, we also confirmed that the CR-induced increase in the *Pck2* transcript is reflected in protein levels (Figures 4h and 4i; SAAR/CD: 5-10 fold, p < 0.05 or lower) and that the changes in Pepck-M are CR-dependent (Figure 4i, CR_β_ < 0.01). These data indicate that CR increases the utilization of mitochondrial oxaloacetate to replenish cytosolic phosphoenolpyruvate, the substrate and product of Pepck-M, respectively. Since CR did not change blood glucose levels, we infer that the pathway involved in CR-induced Ser biosynthesis is serinogenesis (Ser biosynthesis from non-glucose substrates) and not the typical phosphorylated pathway.

**Figure 4.**
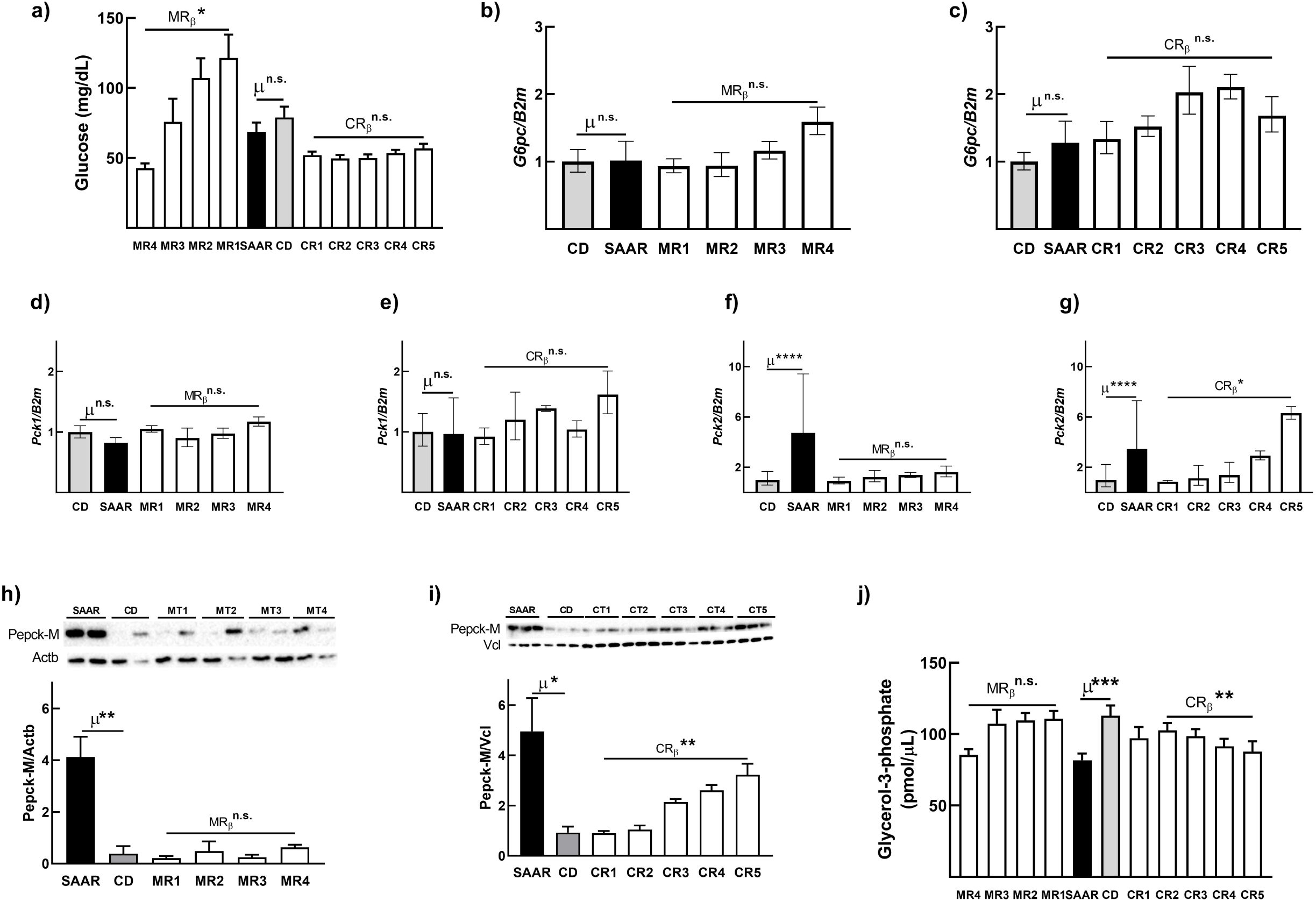

### 5. An increase in *de novo* serine biosynthesis depletes glycerol-3-phosphate required for glyceroneogenesis

The substrates and enzymes of the proximal gluconeogenesis pathway are shared by two other pathways, Ser biosynthesis and glyceroneogenesis. Increased consumption of these substrates by one pathway can affect the end-product concentration of the other. Recent studies demonstrate that an increase in PCK2 expression is associated with an increase in glyceride-glycerol synthesis (Leithner et al., 2018). So, we questioned whether the CR-induced increase in Pepck-M and serinogenesis affects gluconeogenesis and glyceroneogenesis. Although CR upregulated Pepck-M (Figure 4i), it did not alter the mRNA levels of *G6pc* or blood glucose levels. These data suggest that CR is not altering gluconeogenesis. This interpretation is also supported by our previous study in which we demonstrated that mice on the SAAR diet did not change pyruvate tolerance (Ables et al., 2012). To determine whether CR is affecting glyceroneogenesis, we quantified glycerol-3-phosphate (G3P) and found that CR, but not MR, decreases hepatic G3P (Figure 4j, CR_β_ [-0.27(CR2-CR5)] < 0.01). Rats on SAAR had lower hepatic G3P concentrations than those on CD (Figure 4j, SAAR/CD- 0.72, p<0.001). A previous study found that administering ethionine, the ethyl analog of Met, and an inhibitor of Met-dependent functions, including transsulfuration, decreased G3P concentration and thus lipogenesis in both liver and adipose tissue (Tani, Ogata, & Itatsu, 1973). Altogether, these data indicate that CR-induced diversion of 3PG to Ser biosynthesis from glyceroneogenesis, a pathway that provides a major portion of the glyceride-glycerol for re-esterification of fatty acids, is a potential mechanism for CR-specific effects of SAAR on lipid metabolism (Nye, Hanson, & Kalhan, 2008).

### 6. The transcription factor, Nrf2, mediates cysteine restriction-induced serinogenesis

So far, we have investigated the biochemical mechanisms underlying CR-specific effects on hepatic lipid metabolism. We further aimed to discover the molecular mechanisms. Cys is a structural component of the tripeptide glutathione (GSH), an abundant cellular antioxidant. We previously demonstrated that SAAR decreases hepatic non-protein bound total GSH (tGSH = rGSH [GSH in reduced form] + GSSG [GSH in oxidized form]) levels and induces the transcription factor, nuclear factor erythroid 2-related factor 2 (Nrf2) (Nichenametla et al., 2018). Recent studies suggest that Nrf2 regulates Ser biosynthetic enzymes Phgdh, Psat, and Psp, and in concert with another transcription factor Atf4, which induces Pepck-M, regulates stress-related pathways (Chartoumpekis & Kensler, 2013; Chartoumpekis et al., 2018; DeNicola et al., 2015; Kitamura & Motohashi, 2018; Sarcinelli et al., 2020; Shin et al., 2009). Further, it is known that Nrf2 expression depends on dietary fat content, gender, and age (Kerns, Hakim, Zieman, Lu, & Coulombe, 2018; Yin, Corry, Loughran, & Li, 2020; H. Zhang, Davies, & Forman, 2015). We explored whether the GSH/Nrf2/Phgdh/Pepck-M axis is activated in the liver and if its activation is modulated by dietary fat content, gender, and age-at-onset (AAO). We fed eight-week-old male and female C57BL/6J mice with CD and SAAR diets consisting of 10% Kcal or 60% Kcal fat for four months (Supplementary Table 2). In addition, we also fed 18-month-old male and female mice the 10% Kcal fat CD and SAAR diets for three months.

Data from these three models were consistent with our hypothesis. SAAR decreased hepatic GSH, increased Nrf2 protein expression, and translation of its target genes, *Phgdh* and *Pck2* (Figures 5a – 5l). The effect size of SAAR on these markers was dependent on the experimental model. In young male mice, the effect size was, in general, greater on 60% fat than on 10% fat (Figure 5, column 1). In young females, all molecular markers changed in the same direction as observed in the male mice, but changes were primarily similar regardless of the dietary fat content, except for Pepck-M (Figure 5, column 2). With regards to sexual dimorphism, on 10% fat diets, no differences were found between young male and young female mice; however, the effect sizes were significantly larger in males feeding on a 60% fat diet compared to females on a 60% fat diet (Figure 5, column 3, Supplementary Table 3). In males, similar differences were found regardless of AAO, but in females, young-onset had a greater effect size compared to the adult-onset (Supplementary Table 4).

**Figure 5.**
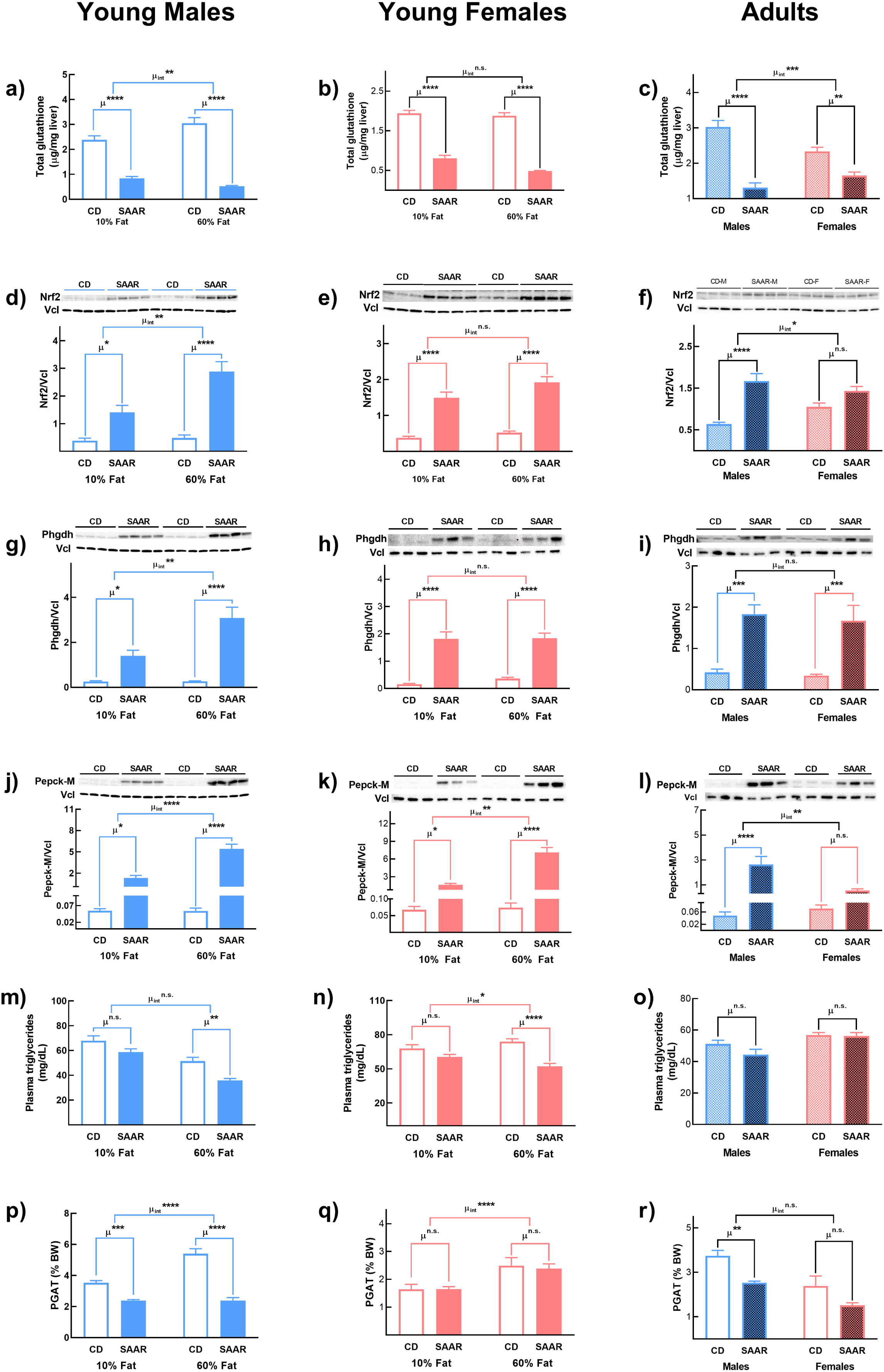

We further explored if the SAAR-induced changes in the molecular markers result in similar changes in plasma triglycerides and adipose depot weights. Overall, there is an agreement between the molecular markers and the ultimate phenotypes, i.e., triglycerides and adipose depot weights in males. Plasma triglycerides were significantly lower in young male, and female mice fed SAAR on a 60% fat diet but not on a 10% fat diet (Figures 5m and 5n). Triglyceride concentrations between CD and SAAR were not different in adult mice (Figure 5o). Perigonadal adipose tissue depot weights (PGAT) were significantly lower in young and adult male mice regardless of the dietary fat content (Figures 5p and 5r) but not in female mice (Figure 5q). Multiple studies report that SAAR-induced responses in adipose metabolism are dependent on gender, AAO, and dietary fat concentration, but the mechanisms are unclear (Forney et al., 2020; Mattocks, Malloy, & Nichenametla, 2019; Nichenametla et al., 2020). Our data suggest that the GSH/Nrf2/Phgdh/Pepck-M axis is an underlying molecular mechanism for CR-specific effects on adipose metabolism. The extent of activation of this pathway determines the magnitude of the changes in triglycerides and adipose depot weight loss.

### 7. Plasma total cysteine and serine, but not methionine, correlate with triglyceride concentrations and metabolic syndrome criteria in humans

To investigate the relevance of our findings from animal models to humans, we retrospectively analyzed data and biospecimens collected from three human studies. Association of plasma amino acids (Met, tCys [reduced Cys + oxidized Cys-Cys + protein-bound Cys], Ser, and tCys/Ser) with triglycerides and the occurrence of MetS criteria were investigated in a subset of individuals from an earlier epidemiological study (Janosikova et al., 2003). To test the effect of dietary fat composition on low SAA diet-induced changes in plasma Ser, we compared Ser concentrations from two short-term controlled-feeding studies (study 1 and study 2), in which high and low SAA diets were formulated with either the same (study 1) or different concentrations of saturated and polyunsaturated fatty acids (study 2). Ser data from one of these studies were previously published (Olsen et al., 2021).

Descriptive statistics of controls and patients with coronary artery disease (CAD) are presented in Supplementary Table 5. Regression estimates and p-values for associations of individual amino acids with triglycerides and MetS criteria are given in Figures 6a-6h and Supplementary Table 6. In the fully adjusted models (adjusted for age, gender, and body mass index), triglycerides did not change with an increase in plasma Met (Figure 6a) but increased by 0.54% (95 % CI: from 0.12 to 0.96, p<0.05, Figure 6b) with a 1% increase in plasma tCys and decreased by 0.30% (95% CI: from - 0.49 to -0.10, p<0.05) with a 1% increase in plasma Ser (Figure 6c). Triglycerides increased by 0.33% with a 1% increase in the ratio of tCys to Ser (95% CI: from 0.15 to 0.52, p<0.05, Figure 6d).

**Figure 6.**
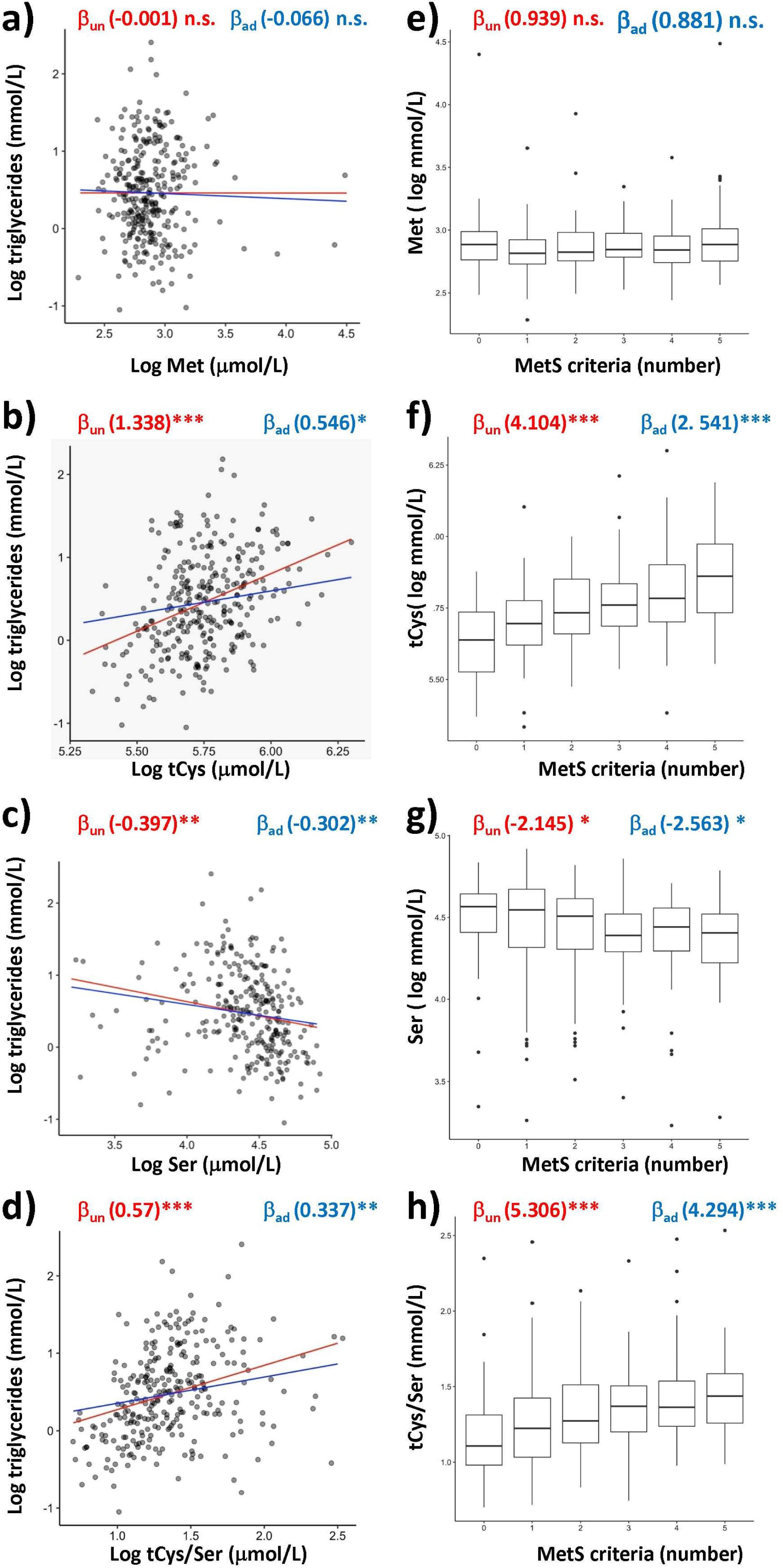

Next, we questioned if plasma Met, tCys, Ser, and tCys/Ser were associated with the occurrence of MetS criteria. Plasma Met was not associated with MetS criteria (Figure 6e). In the unadjusted model, plasma tCys concentration increased by 4.10% (95% CI: from 3.14 to 5.07, p<0.001) with an increase in each MetS criteria (Figure 6f). The estimate for tCys was attenuated but remained significant after adjusting for age and sex (2.54%, 95% CI: from 1.18 to 3.13, p<0.001). In contrast, plasma Ser decreased with an increase in the number of MetS criteria in both models (unadjusted model, by -2.14%, 95%CI: from -4.12 to 0.12, p=0.03, Figure 6g; adjusted model, by - 2.56%, 95% CI: from -4.27 to -0.34, p=0.02); however, the association of tCys/Ser with MetS criteria was stronger than the association with either amino acids alone. In the unadjusted model, the ratio of tCys/Ser increased by 5.31 % with an increase in each MetS criteria (95% CI: from 3.08 to 7.57, p<0.001, Figure 6h). After adjusting for age and gender, the tCys/Ser ratio increased by 4.29%, with increases in each MetS criteria (95% CI: from 1.86 to 6.78, p<0.01). Current literature provides evidence for a strong association of tCys, and a few studies show a modest association of Ser with plasma triglycerides and/or MetS (Fridman et al., 2021; Mook-Kanamori et al., 2016). Our data show that the ratio of tCys/Ser might serve as a better biomarker for MetS than each of them individually and is consistent with mechanisms suggested by our animal data, i.e., tCys deficiency results in higher Ser biosynthesis and lower triglycerides (Figures 6f and 6h).

### 8. Low-SAA diets increase plasma serine depending on the dietary fatty acid composition

We show that our epidemiological findings are clinically relevant by presenting data from two previously conducted short-term controlled feeding studies. i.e., low SAA diets increase plasma Ser depending on the dietary fat composition (Olsen et al., 2020; Olsen et al., 2018, 2021). In Study 1, participants were fed diets with either low (SAA_low_) or high (SAA_high_) SAA (Supplementary Table 7). In Study 2, the type of fat in both diets was changed to impact triglyceride synthesis, i.e., SAA_low_ and SAA_high_ diets were supplemented with high polyunsaturated fatty acids (SAA_low+PUFA_) and high saturated fatty acids (SAA_high+SFA_), respectively. Both studies were conducted for one week, and changes in plasma Ser on days 0, 3, and 7 were compared between the diet groups. Dietary SAA concentration, fatty acid concentrations, and sample sizes are provided in Supplementary Table 7.

Compared to plasma Ser on day 0, the estimated marginal mean Ser concentration on day seven slightly decreased in both SAA_low_ (from 100.8 to 96.8 mmol/L) and SAA_high_ (from 103.7 to 100.7 mmol/L) groups in Study 1. The time-related changes between the two diet groups were similar (Figure 6i and Supplementary Table 8, p_int_ = 0.94). In Study 2, there was an increase in estimated marginal mean plasma Ser concentrations from 106 to 118 mmol/L in the SAA_low+PUFA_ group and a decrease from 101.2 to 96.1 in the SAA_high+SFA_ group, and these time-related changes between the two diet groups were significantly different (Figure 7 and Supplementary Table 8, p_int_<0.05). The effect of high dietary SAA concentration in the presence of saturated fat, which promotes triglyceride synthesis, on decreasing the plasma Ser is comparable to the findings from mouse models, i.e., stronger effects of SAAR diet on Phgdh and Pepck-M protein expression on 60% fat diet, which increases fatty acid/triglyceride cycling, than on 10% fat diet (Figures 5g, 5h, 5j, and 5k). Collectively, the associations between SAA, Ser, triglycerides, and MetS criteria in human studies are consistent with mechanistic findings from animal models.

**Figure 7.**
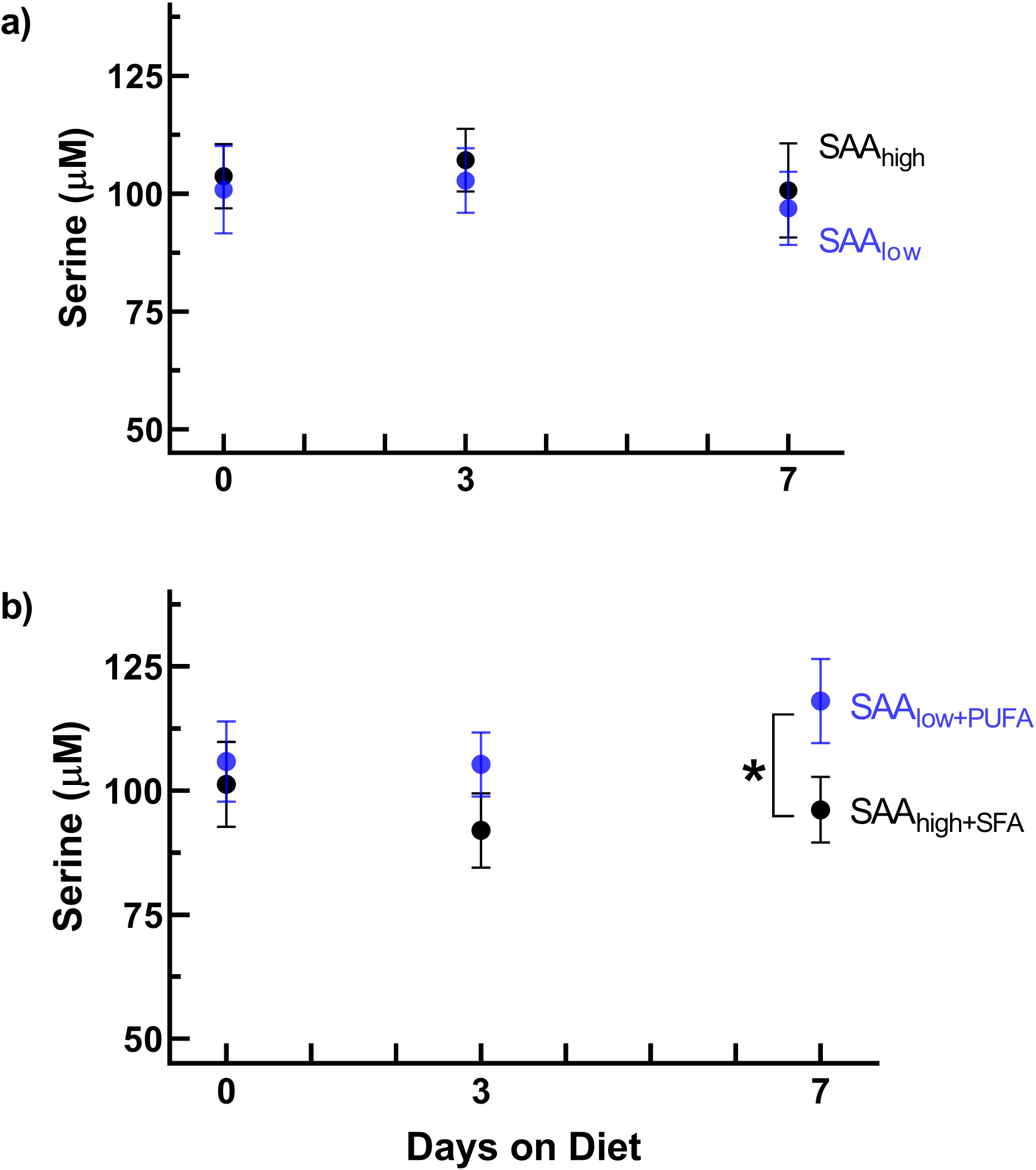

## Discussion

A salient feature of SAAR is the amelioration of lipid metabolism homeostasis, but its mechanisms are ill-defined. To understand the molecular mechanisms, we conducted dietary SAA depletion-repletion bioassays. Data show that MR and CR exert discrete and non-nutritive biological effects. MR-dependent, SAAR-induced changes include decreases in growth rate, blood glucose, plasma Igf1, leptin, increases in food consumption, and Fgf21 (Figures 1 and 4a). CR-dependent SAAR-induced changes include increases in plasma adiponectin, Ser, Thr, and His; decreases in plasma Met, Trp, and Phe (Figures 1 and 2). Except for a few plasma amino acids, most of the markers altered by SAAR responded to the dose of either MR or CR. We propose a mechanism for CR-specific effects on lipid metabolism based on the molecular data. The SAAR-induced decrease in hepatic tGSH induces Nrf-2 expression and its downstream effectors, Phgdh and Pepck-M, which increase serinogenesis. Higher serinogenesis depletes the substrates required for glyceroneogenesis, which provides most of the glyceride-glycerol to re-esterify fatty acids, eventually resulting in lower adipose depot weights. We also show that the SAAR-induced changes in molecular markers are consistent and reproducible in different animal models. However, the magnitude of changes depend on AAO (young > adult), gender (male > female), and dietary fat concentration (60% fat > 10% fat). Consistent with the mechanistic data from animal models, we found a negative correlation between plasma Ser and plasma triglycerides and MetS criteria in humans, while plasma Cys positively correlated with both; plasma Met did not show any correlation. Additionally, we establish the clinical relevance of our findings by showing that plasma Ser is amenable to SAAR-like diets in humans when combined with high dietary intakes of polyunsaturated fatty acids.

Our current understanding of the roles Cys plays in energy metabolism is relatively less than that of its functions associated with thiol-disulfide exchanges. Previous reports show that Cys can affect energy metabolism by contributing its carbon moiety to biomass and as a signaling molecule. In pancreatic cancer cell lines, Cys catabolism contributed up to 10% of intracellular pyruvate, and this contribution increased to 20% upon knocking out pyruvate kinase (L. Yu et al., 2019). Other reports show that Cys can affect energy metabolism through mTOR signaling (Bonifacio, Pereira, Serpa, & Vicente, 2021; Gu et al., 2017). Our data show that Cys plays a much broader role in regulating central carbon metabolism than merely contributing to its carbon skeleton. By depleting GSH, CR increased the expression of Pepck-M, which can alter TCA flux. Phgdh, the second CR-induced enzyme, is typically associated with glycolytic flux (Locasale et al., 2011). However, our data show that it can alter glyceroneogenesis as well. Our study also indicates that through its thiol-disulfide function, Cys can affect adipose metabolism through adiponectin secretion, a marker well-associated with energy metabolism (Lee & Shao, 2014). We did not investigate the mechanisms of CR-induced adiponectin secretion. But, previous *in vitro* studies demonstrate that intracellular retention and secretion of adiponectin are dependent on thiol concentrations (Wang et al., 2007). We speculate that the increased adiponectin secretion in our study could be due to lower glutathione concentration in the adipose tissue. Overall, our data provide a mechanistic basis for the effects of Cys on metabolic health. From the perspective of disease prevention, our data also emphasize the importance of considering the individual concentrations of dietary Met and Cys and their effect on energy metabolism, mainly when the total SAA content is low. Three diets with different ratios of Cys to Met (0.2, 1, and 2), despite all having the same total SAA concentration, had different effects on body weight, blood urea nitrogen, and bile acids in rats (Sarwar, Peace, & Botting, 1991). The nutritional requirements of Met and Cys, both for laboratory animals and humans, are always considered together. The above findings warrant consideration of the ratio of Met to Cys in addition to total SAA concentrations in evaluating the effects of SAAR on human health and diseases.

Due to its involvement in one-carbon metabolism, there is considerable interest in treating cancer by inhibiting Ser biosynthesis (Amelio, Cutruzzola, Antonov, Agostini, & Melino, 2014). Contrarily, only a few studies investigated the role of Ser on altered lipid profiles. Whether Ser affects body adiposity, and if so, the mechanisms associated are not well-studied (Muthusamy et al., 2020). Our data suggest that serinogenesis lowers glyceroneogenesis by limiting the availability of G3P (Figure 4j). Other studies prove the corollary, i.e., inhibition of Phgdh deceased plasma Ser and simultaneously increased hepatic fat content in mice fed Met-choline deficient diets (Sim et al., 2020). Of note, the composition of the Met-choline deficient diet is different from that of the SAAR diet. The former is deficient in both methyl donors, i.e., Met and choline; hepatic steatosis occurs due to low VLDL secretion (Marcolin et al., 2011). Although the SAAR diet is low in Met, it is not deficient in methyl donors due to abundant choline. Unlike Met-choline deficient diets, it decreases hepatic steatosis (Malloy et al., 2013). In addition, Phgdh transgenic mice on a high-fat diet had lower hepatic triglycerides than the wild-type mice, indicating that an increase in Ser biosynthesis, independent of the CR diet, can affect lipid metabolism (Sim et al., 2020). The epidemiological association of *PHGDH* DNA methylation with triglyceride levels suggests that Ser might play a similar role in lipid metabolism in humans and animals (Truong et al., 2017). This finding is consistent with results from our epidemiological study where plasma Ser was positively associated with triglycerides and MetS criteria (Figure 6). Another epidemiological study shows that diabetic patients had significantly lower Ser than controls (Y. Zhou et al., 2013). Considering all this data, we propose that serinogenesis could be an excellent target for treating hypertriglyceridemia and other diseases associated with lipid metabolic dysregulation.

Our data suggest lower body adiposity could be a consequence of increased serinogenesis but do not explain whether increased Ser availability has any functional implications. Similar to lifespan extension and higher Ser concentrations induced by the SAAR diet in rodents, studies in yeast suggest that Ser supplementation increases lifespan, probably by supporting one-carbon metabolism (Enriquez-Hesles et al., 2021). However, specific molecular mechanisms remain unclear. Although Ser can support one-carbon metabolism as a methyl donor, it is less likely that animals increase serinogenesis for methyl groups as the SAAR diet has other fungible methyl donors - choline and glycine (Supplementary Table 1a). Ser can also increase endogenous Cys biosynthesis by condensing with homocysteine. Since the SAAR diet lacks Cys, higher Ser biosynthesis to support Cys biosynthesis might seem plausible. However, it is also improbable, as Ser contributes only to the carbon skeleton in Cys; the S moiety comes from Met, which is limited in the diet (du Vigneaud, Kilmer, Rachele, & Cohn, 1944). Hence, higher Ser availability alone cannot increase Cys biosynthesis in SAAR animals. After obtaining a methyl group from Ser in the one-carbon cycle, tetrahydrofolate is converted to 5,10-methylene tetrahydrofolate, which reduces NADP to NADPH. Thus Ser indirectly contributes to the generation of reducing equivalents (NADPH), which are required for multiple purposes. One recent report shows that Ser supports hepatic fatty acid synthesis by providing NADPH (Z. Zhang et al., 2021). Previous studies show that SAAR induces a futile lipid cycle, i.e., an increase in lipogenesis and lipolysis (Perrone et al., 2008). Thus, a higher serinogenesis might provide NADPH required for fatty acid biosynthesis.

A second functional implication of higher Ser availability and its ability to increase NADPH is the maintenance of the redox state of GSH, which is involved in a multitude of cellular functions, including growth, stress response, and detoxification of reactive oxygen species (Forman, Zhang, & Rinna, 2009). Healthy tissues in well-fed animals have abundant tGSH and maintain a high ratio of rGSH to GSSG (≥10) (Kosower & Kosower, 1978). Maintenance of this ratio is essential to carry out normal cellular functions, and its decrease is associated with several diseases (Choromanska et al., 2020). A decrease in hepatic tGSH is a characteristic feature of SAAR (Nichenametla et al., 2018; Richie et al., 2004). The only way to sustain GSH-dependent redox processes is by increasing the turnover of its redox cycling (GSH ↔ GSSG), which requires NADPH as a cofactor, more so in mitochondria than in cytoplasm due to the production of reactive oxygen species. Since Ser can enter mitochondria and contribute to the reduction of NADP^+^ to NADPH through 5,10-methylenetetrahydrofolate, we speculate that SAAR induces serinogenesis to maintain the mitochondrial milieu in a reduced state and that its effect on glyceroneogenesis is a collateral health benefit. A previous study supports our explanation which shows that Pepck-M, the mitochondrial enzyme induced by SAAR, was critical in maintaining GSH pools in a reduced state (Bluemel et al., 2021). Our hypothesis is also supported by another study in which mice were depleted of GSH by feeding buthionine sulfoximine, an inhibitor of the GSH biosynthetic enzyme, Gclc (Elshorbagy et al., 2016). Despite a decrease in hepatic non-protein-bound total GSH concentration, these mice had similar levels of rGSH and exhibited a lean phenotype; however, this study did not quantify Phgdh and Pepck-M. We also speculate that the higher ratio of rGSH to tGSH might increase mitochondrial fidelity and contribute to lifespan extension, the classical feature of the SAAR diet.

A unique feature of SAAR-induced serinogenesis is that the oxaloacetate is sourced from mitochondria, not from the cytosol, i.e., an increase in the expression of Pepck-M, but not Pepck-C. Both Pepck-C and Pepck-M can alter gluconeogenesis flux and Ser biosynthesis (Lowry et al., 1987). However, the preferential upregulation of Pepck-M by SAAR remains unclear. One possible explanation might be the presence of an amino acid response element in the *Pck2* promoter while, to our knowledge, no such promoter has been reported for *Pck1* (Mendez-Lucas, Hyrossova, Novellasdemunt, Vinals, & Perales, 2014). It was also reported that Atf4, another transcription factor, binds to the response element in *Pck2* to regulate amino acid metabolism (S. Yu, Meng, Xiang, & Ma, 2021). We and others previously demonstrated that unfolded protein response and the Perk-eIf2α-Atf4 pathway, which are activated by amino acid limitation, are also upregulated by the SAAR diet (Mendez-Lucas et al., 2014; Nichenametla et al., 2018; Pettit et al., 2017). Altogether, these lines of evidence suggest that the preferential upregulation of Pepck-M might be occurring to maintain mitochondrial protein homeostasis. This idea is consistent with the fact that, despite the lower cytosolic protein synthesis, the SAAR diet did not alter the mitochondrial protein synthesis rates (Nichenametla et al., 2018; Pettit et al., 2017). In addition to maintaining mitochondrial protein synthesis, an anabolic process, SAAR-induced serinogenesis might also contribute to the biosynthesis of nucleotides required for cellular proliferation, another anabolic process, suggesting that SAAR-induced changes in central carbon metabolism are directed toward biomass accumulation, i.e., growth. However, maintaining these biosynthetic processes depletes oxaloacetate from the TCA cycle, lowering the NADH production and thus ATP yield. The ability of living systems to delicately balance the utilization of carbon and nitrogen sources between biomass accumulation (gluconeogenesis, nucleotide and protein synthesis) and energy production (ATP yield) under environmental constraints such as nutrient limitation were recognized earlier and is a typical trade-off observed among life-history traits (Burger, Hou, & Brown, 2019; Chen & Nielsen, 2019). However, despite having *ad libitum* access and similar total amino acid content in SAAR, CR-titrated diets, and CD, such adaptation is quite interesting. The only difference in these diets was the Cys concentrations, in other words, the availability of thiol groups. Changes in energy metabolism at the organismal level based on the thiol-group abundance may not seem plausible. Yet, such adaptations occur in unicellular organisms such as yeast, where tRNA thiolation alters the carbon and nitrogen homeostasis. Mutant tRNAs that cannot be thiolated resulted in an amino acid starvation phenotype despite sufficient availability of amino acids and switched metabolism from nucleotide synthesis to the synthesis of trehalose, a storage carbohydrate (Gupta et al., 2019). Overall, we propose that higher Pepck-M induced by CR rewires the metabolic pathways to sustain growth and survival under the limited availability of organic unsubstituted thiol (SH) groups at the cost of mitochondrial respiration.

Our data demonstrate that the SAAR-induced phenotype is a combination of MR and CR and that SAAR induces metabolic flexibility by upregulating two enzymes, Phgdh and Pepck-M. Based on epidemiological data, we believe that pharmacological upregulation of these two enzymes might prove beneficial in treating obesity, metabolic syndrome, and associated diseases. Alternatively, pharmacological CR might also induce serinogenesis. Indeed, unpublished data from our group show a successful and well-tolerated lowering of plasma tCys using a single dose of MESNA (2-mercaptoethan sulfonate sodium) in overweight or obese men without affecting Met concentrations (Vinknes, 2020). In addition, the SAAR diet may be used as adjuvant therapy in applicable conditions. In diabetic patients, gluconeogenesis becomes less sensitive to the prandial fluctuations and remains a constitutive source of blood glucose (Gastaldelli et al., 2000; Wajngot et al., 2001). PPAR-γ agonists, such as rosiglitazone, improve blood glucose homeostasis by inhibiting gluconeogenesis and inducing glyceroneogenesis in adipose tissue (Cadoudal et al., 2007; Gastaldelli et al., 2006; Leroyer et al., 2006). While these dual effects make rosiglitazone an effective anti- hyperglycemic agent, it promotes weight gain, is less efficient in individuals with higher body mass index, and increases the risk of myocardial infarction (Fonseca, 2003; Leroyer et al., 2006; Monami, Marchionni, & Mannucci, 2008). Since SAAR-induced serinogenesis depletes the proximal substrates from glyceroneogenesis, decreases blood glucose levels, and induces weight loss simultaneously, it would be a better option than the low-calorie diets currently used for weight-management programs for patients receiving thiazolidine treatment (Asnani, Richard, Desouza, & Fonseca, 2003).

Our current study has some limitations. SAAR-induced changes in adult mice were milder than changes observed in young mice. Nutritional requirements for growing rodents are much higher than for adult and old rodents (Ishibashi & Kametaka, 1977). It remains unknown whether the decrease in growth-related demands for SAA would differently affect the partitioning of Met into Met and Cys in post-growth life stages. Future studies should explore whether CR- and MR-titrated diets induce similar changes in adult and old animals as in young rodents by further decreasing the concentration of Met in SAAR diets, i.e., below 0.12% Met. Our conclusions on SAAR-induced changes, i.e., increase in serinogenesis and decrease in glyceroneogenesis, are based on transcriptional and translational changes in proteins and metabolites involved in these pathways. Tracer-based studies are required to confirm our findings and determine whether SAAR alters the flux of these two pathways. Although conclusions from our epidemiological analysis are congruent with the mechanisms observed in animals, we cannot establish causality. Long-term feeding studies with SAAR-like diets using metabolic tracers will help determine the causality in humans.

## Methods

### Animal studies design

Animal experiments were conducted in five different cohorts, two cohorts of rats and three cohorts of mice.

#### Experiment 1

Discrete effects of MR were determined by feeding six groups of eight-week-old male F344 rats (n = 8/group) for 12 weeks. The diets were: CD diet (0.86% w/w Met without Cys), SAAR diet (0.17% w/w Met without Cys), and four MR diets, all replete with 0.5% w/w Cys but with progressively restricted Met (MR1 - 0.17%, MR2 - 0.10%, MR3 - 0.07%, and MR4 - 0.05% w/w).

#### Experiment 2

Discrete effects of CR were determined by feeding the second cohort of seven groups of eight-week-old male F344 rats (n = 8/group) for 12 weeks. The diets were: CD diet, SAAR diet and five more diets, all with 0.07% w/w Met but with progressively restricted Cys (CR1 - 0.5%, CR2 - 0.25%, CR3 - 0.12%, CR4 - 0.06%, and CR5 - 0.03% w/w).

#### Experiment 3

Effects of dietary fat content on SAAR-induced changes were determined by feeding the first cohort of eight-week-old male C57BL/6J mice CD and SAAR diets containing 10% Kcal and 60% Kcal fat (four groups of mice; n = 8/group) for 16 weeks.

#### Experiment 4

To investigate gender-specific effects, the second cohort of eight-week-old C57BL/6J female mice were fed similar diets as male mice (four groups of mice; n = 8/group) for 16 weeks.

#### Experiment 5

AAO-dependent effects were determined by feeding the third cohort of 18-month-old male and female C57BL/6J mice, CD and SAAR diets with 10% Kcal fat (four groups of mice; n = 8/group) for 12 weeks.

#### Animal procedures

All animal procedures were conducted according to the guidelines of the Institutional Animal Care and Use Committee of Orentreich Foundation for the Advancement of Science. Throughout the study period, all animals were singly housed and provided with nestlets (Ancare, Corp., Bellmore, NY) for enrichment. Food (chow and experimental diets) and acidified water were provided *ad libitum*. All animals were maintained on a 12-hr: 12-hr light-dark cycle, with room temperatures of 20±2°C and 50±10% relative humidity. Food intake and body weights were recorded weekly.

For experiment 1, four-week-old male F344 rats were obtained from Charles River Laboratories (Wilmington, MA, strain # 002), quarantined for one week, and provided with Laboratory Rodent Diet 5001 (PMI Nutrition International, Brentwood, MO) and acidified water *ad libitum* until they reached eight-weeks of age. Rats were then randomly divided into six groups and fed the experimental diets. After 12 weeks on the diet and following an overnight fast, the rats were anesthetized with isoflurane (Piramal Critical Care, Bethlehem, PA, USA), and blood was collected from the retro-orbital plexus into EDTA-Vacutainer tubes (BD Biosciences, Franklin Lakes, NJ). The blood tubes were kept on ice centrifuged at 15000 g for 15 min at 4°C; the plasma was flash-frozen in liquid nitrogen and then stored at -80°C. Rats were then sacrificed by CO_2_ asphyxiation followed by decapitation. Livers were weighed, snap-frozen in liquid nitrogen, and stored at -80°C. An aliquot of the liver was incubated overnight in RNA*later* (ThermoFisher Scientific, AM7021, Waltham, MA) at 4°C. The next day, RNA*later* was discarded, and tissues were stored at -80°C until processed for RNA. Upon analysis of the data from the first study, the second cohort of eight-week-old rats was obtained. Animal husbandry and experimental procedures were similar to those followed for the first cohort; however, the second cohort was fed CD, SAAR, CR1, CR2, CR3, CR4, and CR5 diets.

Male and female C57BL/6J mice were obtained from the Jackson Laboratory (stock # 000664, Bar Harbor, ME) and quarantined for one week. The mice (8 weeks of age) were fed CD and SAAR diets containing either 10% Kcal fat or 60% Kcal fat for 16 weeks. Adult mice were aged in-house and, upon reaching 18-months of age, were fed CD and SAAR diets for 12 weeks. At the study’s conclusion, blood and livers were obtained from 4-hr fasted mice and were processed in the same manner as described in the rat studies.

#### Animal diets

All diets used in the rat cohorts were isocaloric and contained the same amount of total amino acids as the CD diet (Supplementary Tables 1a and 1b, Research Diets, New Brunswick, NJ). The experimental diets were analyzed to verify the amino acid content (Covance, Madison, WI) prior to the start of the study. Considering the losses expected for Cys during analytical procedures, the dietary content of Met and Cys were within the expected range (Supplementary Table 9) (Lamp, Kaltschmitt, & Ludtke, 2018). The other amino acids were also within the expected ranges (data not shown). The diets used for mouse studies were also isocaloric and contained the same amount of total amino acids (Supplementary Table 2). The energy density of CD and SAAR diets with 60% Kcal fat were similar but different from that of 10% Kcal fat CD and SAAR diets; both formulations were used in previous studies (Ables et al., 2012; Cooke et al., 2020). While the Met content of mouse CD diets was the same as the rats’ CD diets, i.e., 0.86%, Met was slightly higher in rat SAAR diets (0.17%) than in mouse SAAR diets (0.12%). The lower Met content in the mice SAAR diet was based on our unpublished experience that SAAR-induced changes attenuate over time if the dietary Met content was 0.17%. All diets were stored at -20^0^C throughout the study period.

### Epidemiological study design and subject selection

Details of the study subject recruitment of the parent study, which investigated the effect of genetic variation in homocysteine metabolizing enzymes on CAD risk, were described elsewhere (Janosikova et al., 2003). In the current cross-sectional study, a subset of individuals (n = 309) was randomly selected from the 278 CAD patients and 591 controls. All subjects were divided into six groups according to the number of MetS, ranging from 0 to 5. MetS criteria were assigned according to Alberti et al., with one modification; due to the unavailability of waist circumference, abdominal obesity was classified according to WHO 1999 criteria based on waist-hip ratio >0.90 in men and >0.85 in women (Alberti, Zimmet, & Shaw, 2006; Lean, Han, & Morrison, 1995). For laboratory analyses of Ser and additional metabolites, we randomly selected a subset of subjects (n=50) from each group with 0, 1, 2, 3, and 4 MetS criteria while maintaining the proportion of controls versus CAD patients and gender, and all 59 subjects meeting all 5 MetS criteria. Sulfur amino acids (Met and tCys), Ser and triglyceride concentrations were quantified in the baseline plasma. Two CAD patients were excluded on the basis of severe hypertriglyceridemia (TAG > 10 mmol/L) and the presence of co-morbidities including pancreatitis and liver disease, leaving a total number of 307 available for analysis. Ser was quantified from 287 subjects only.

### Controlled-feeding study design and dietary formulation

Study subject recruitment details were published previously (Olsen et al., 2020; Olsen et al., 2018, 2021). In brief, two randomized, controlled, double-blind pilot studies were performed from 2016 to 2018 at the Centre for Clinical Nutrition at the Institute of Basic Medical Sciences, University of Oslo, Norway. Study 1, in which only SAA content was altered, was performed in overweight and obese women, whereas Study 2, in which SAA and fatty acid composition were altered, was performed in normal-weight men and women. In both studies, diets were designed following the Nordic Nutrition Recommendations to meet percent energy requirements for protein, carbohydrates, and fats and to be isocaloric across the intervention groups (*Nordic Nutrition Recommendations 2012*□: *Integrating nutrition and physical activity*, 2014). Diets were designed with fixed amounts of SAA per day with the goal of staying around the estimated average requirements in mg/kg body weight/day for each subject. Sample size and dietary compositions for both studies are described in Supplementary Table 7.

#### Amino acid analyses

##### Animal studies

Plasma-free amino acid levels were determined by reverse-phase ultra-performance liquid chromatography and UV detection. Separation chemistry is described in detail elsewhere (Bidlingmeyer, Cohen, & Tarvin, 1984; Heinrikson & Meredith, 1984). Briefly, plasma was deproteinized with an equal volume of 10% sulfosalicylic acid containing an internal standard (Norvaline, Catalog No. PHR2210, Sigma-Aldrich, St Louis, MO). Samples were centrifuged at 10000 g for 5 min at room temperature, and supernatants were derivatized with AccQTag Ultra derivatization kit according to the manufacturer’s instructions (Waters Corporation, Milford, MA, USA). Derivatized amino acids were separated on a C-18 column (2.1 mm × 100 mm, particle size = 1.7 μm) maintained at 55 °C (Waters Corporation, Milford, MA, USA) with UV detection (260 nm). Concentrations were determined by comparing peak areas with known standards using Waters Empower 2 Software and expressed in μM (Waters Corporation, Milford, MA, USA). To ensure no drift, standards were run between every ten samples.

Rat hepatic total Ser was quantified using a fluorometric DL-Serine Assay Kit (catalog No. K743, Biovision, Milpitas, CA, USA). The assay was based on enzymatic conversion of L-Ser to D-Ser, which is converted to an intermediate product. The intermediate is further oxidized and converted to a stable fluorophore. Briefly, frozen liver aliquots were homogenized in Serine Assay Buffer in a Potter-Elvehjem homogenizer and centrifuged at 15000 g, for 10 min at 4°C. The supernatants were deproteinized with Sample Cleanup Mix and filtered through 10 kDa molecular weight cutoff spin columns (Corning, catalog 431478). Filtrates were used in the assay, and all procedures recommended by the manufacturer were followed. Background fluorescence for each sample was quantified without the addition of the enzyme. Net fluorescence of the samples was extrapolated to a standard curve. Values were expressed as nmol/mg tissue.

##### Epidemiological cohort

Plasma tCys was quantified as described earlier (Krijt, Vackova, & Kozich, 2001). Met and Ser in plasma were determined by LC-MS/MS using a commercially available kit for amino acid analysis (EZ:faast™, Phenomenex, Torrance, USA). The internal standard solution, which is a component of the kit, was supplemented with 100 μmol/L of serine-d_3_ (Cambridge Isotope Laboratories Inc., USA). The LC-MS/MS system consisted of the Agilent 1290 Infinity LC System (Agilent Technologies, Palo Alto, CA, USA) coupled with an API 4000 triple quadrupole mass spectrometer (Applied Biosystems, Foster City, CA, USA). All plasma amino acid concentrations were expressed as μM.

##### Short-term human feeding studies

Plasma Ser concentrations from both studies were measured with LC-MS/MS as reported previously (Antoniades et al., 2009; Olsen et al., 2020; Olsen et al., 2021). All plasma amino acid concentrations were expressed as μM.

#### Hepatic glycerol-3-phosphate

G3P was assayed in rat livers using a fluorometric kit (Catalog No. K196, PicoProbe Glycerol-3-Phosphate Assay Kit, BioVision, Milpitas, CA, USA) following the manufacturer’s recommendation. Small aliquots (20 mg) of livers were homogenized in a 200 ul ice-cold assay buffer. The lysate was centrifuged at 10,000 g for 5 minutes at 4°C, and the supernatant was transferred to a ten kDa molecular weight cut-off spin column. The columns were spun at 12000 g for 30 minutes at 4°C, and the resultant filtrate was stored at -80°C until assayed. The standards, samples, and background controls were assayed in duplicate. A complete reaction mixture (50 uL per well) was added to the standards and samples, and a background control was run for each sample by adding 50 ul of a mixture comprised of all reagents except the Enzyme Mix to those wells. Fluorescence was measured on a plate reader (Molecular Devices, San Jose, CA, USA) at excitation/emission wavelengths of 535/587 nm, respectively. Background-corrected sample fluorescence was compared to the standard curve to obtain the G3P content of the samples. Although enough filtrate was obtained to perform the assay, some were retained in the spin columns. Since we were not able to collect and quantify all the filtrate from the spin column, concentrations are expressed as the pmol/ul filtrate rather than pmol/mg liver.

#### Glutathione

Non-protein-bound tGSH in the liver was determined by an enzymatic recycling method using Ellman’s reagent (Tietze, 1969). Briefly, aliquots of the liver were homogenized in four volumes of 5% metaphosphoric acid and incubated on ice for 20 min. Supernatants obtained from the lysates by centrifugation at 15000 x g/20 min/4°C were further diluted before using in colorimetry. GSH levels were expressed as mM.

#### Transcriptional changes

Transcriptional changes were determined by quantifying the mRNA expression of genes of interest relative to the house-keeping gene, β2-microglobulin. Total RNA was extracted from RNAlater-treated livers using TRIzol Reagent (Catalog No. 15596026, ThermoFisher Scientific, Waltham, MA, USA) following the manufacturer’s recommendations. RNA pellets were dissolved in RNase-and-DNase-free water. Purity and concentration were determined based on the absorbance ratio at 260/280 nm using NanoDrop (ThermoFisher Scientific, Waltham, MA). Before transcribing the mRNA to cDNA using the High Capacity cDNA Reverse Transcription Kit (Catalog No. 4368813, Applied Biosystems, Waltham, MA USA), residual DNA was digested with DNase I (Catalog No.18068015, Invitrogen, Waltham, MA). The cDNA obtained was diluted 10- fold with DNase-and-RNase-free water and used for real-time TaqMan PCR. Inventoried assays (Supplementary Table 10b, ThermoFisher Scientific, Waltham, MA) for target genes were selected such that the primers span exon-exon junctions. β-2 Microglobulin was used as the housekeeping gene for all reactions, and fold-change in mRNA expression of each diet group was expressed after normalizing with that of the CD group.

#### Translational changes

##### ELISAs

Changes in protein expression were determined by using ELISA (plasma proteins) and semi-quantitative western blots (tissue proteins). Growth factors (Igf1 and Fgf21) and adipokines (leptin and adiponectin) were determined by quantitative sandwich ELISAs following the manufacturer’s recommendation (Supplementary Table 10a, Quantikine ELISA kits, R&D Systems Inc., Minneapolis, MN, USA) and expressed either as pg/mg or ng/mg of plasma protein. Samples were always run in duplicate. Depending on the standard curve range and estimated concentrations of analytes in each diet group, plasma was appropriately diluted. If samples were run in multiple plates, a pooled sample was run in all plates.

##### Western blots

Tissues were homogenized with Potter-Elvehjem homogenizers in ice-cold RIPA buffer (50 mM Tris-HCl pH 8.0, 150 mM NaCl, 1% Triton X-100, 0.5% Sodium Deoxycholate, 0.1% SDS) containing protease and phosphatase inhibitors (Catalog No. 5872, Cell Signaling, Danvers, MA). The lysate obtained was sonicated and incubated on ice for 30 minutes. They were centrifuged at 10000 g/10 min/4°C, supernatants were separated and recentrifuged at 15000 g/10 min/4°C, and the final supernatants obtained were frozen until used for Western blotting. Protein concentration was quantified by a BCA assay (Catalog No. 23225, ThermoFisher Scientific, Waltham, MA). Before loading onto the wells, lysates were denatured by mixing with Laemmli Sample Buffer containing mercaptoethanol and boiling for 5 min. Samples were electrophoresed in either Mini-PROTEAN^R^ or Criterion™ TGX™ precast polyacrylamide gels at 200 V (constant voltage). Proteins were transferred to PVDF membranes using Mini Trans-Blot Cells or a Criterion™ Blotter for 1 hour at 100 V (constant voltage) or with a SureLock™ Tandem Midi Blot Module for 30 minutes @ 25V (constant voltage). The PVDF Membranes were blocked with 5% BSA (Cell Signaling Technology Inc., Danvers, MA) or 5% nonfat dry milk (Bio-Rad Laboratories, Inc.) in 1X TBS containing 0.1% Tween 20 for 1 hour at room temperature. They were then incubated overnight at 4 °C with primary antibodies at dilutions recommended by the manufacturers. After overnight incubation, the membranes were washed three times with TBST and incubated with secondary antibodies for 1 hour at room temperature. The washes were repeated, and membranes were incubated with either Clarity™ Western ECL Blotting Substrate (Bio-Rad) or SuperSignal™ West Femto Maximum Sensitivity Substrate (ThermoFisher Scientific). Bands were visualized using the ChemiDoc XRS+ system (Bio-Rad Laboratories, Inc.). Either β-actin or vinculin was used as the loading controls. Band intensities for all proteins of interest were determined using Image Lab software (Bio-Rad Laboratories, Inc.) and normalized to the intensity of bands from loading controls. To account for technical variation between gels, a batch control with pooled total protein was run on all gels. The fold change in target proteins was expressed as a relative ratio to the loading controls. The sample size for each diet group ranged from 4-8. Sometimes membranes were reused to quantify a second target protein. In this case, membranes were stripped by using Restore™ PLUS Western Blot Stripping Buffer (Catalog No. 46430, ThermoFisher Scientific, Waltham, MA) and incubating them at 37°C for 15 min. Antibody sources and specific conditions used for each antibody are presented in Supplementary Table 10.

##### Plasma Glucose

Fasting blood glucose (16-hr) was determined after decapitation using FreeStyle Lite^®^ glucometers (Abbott Laboratories, Abbott Park, IL, USA) and expressed as mg/dL.

##### Plasma triglycerides

Plasma triglycerides were quantified in microplates using Infinity Triglycerides Liquid Stable Reagent (ThermoFisher Scientific, Waltham, MA, Cat # TR22421) and a Triglycerides Standard from Pointe Scientific (T7531-STD). Triglycerides Reagent (200 ul) was added to wells containing 10 uL plasma and incubated at 37°C for five minutes. The absorbance at 500 nm with a reference wavelength of 660 nm was measured, and the values were extrapolated to a standard curve. Triglyceride concentrations were expressed as mg/dL.

## Statistics

Mean differences between CD and SAAR in the two rat cohorts were assessed by using the two-tailed Student’s t-test. For quantitative data such as concentrations of plasma analytes, CD and SAAR groups from the two cohorts were combined to increase the statistical power and for easier graphical representation; however, semi-quantitative data such as mRNA and protein expression from the two cohorts were tested individually and represented in two different panels. Data were analyzed in R (v 4.1.0) and GraphPad Prism. A p-value of 0.05 was considered statistically significant in all analyses. Further statistical tests to detect the dose-response relationship in MR and CR diets were done using linear mixed-effect regression models and trend tests.

### Linear mixed effect model

Linear mixed-effect regression to test if there were any significant differences in the mean for each of the MR and the CR groups with respect to SAAR. Separate regressions were run for each of the phases. A random intercept was added to models to consider the heterogeneous effects contributed by each sample. All the p-values were adjusted for multiple comparison error using False Discovery Rate (FDR).

### Trend tests

Two sets of estimated mean differences were extracted from the linear mixed models, namely MR groups vs. SAAR and CR groups vs. SAAR. Assuming a linear trend, two simple linear regressions were fitted for each of the two sets. We tested if there was any significant difference between the two estimated slopes using the Welch two-sample t-test (without the assumption of homogeneity).

### Equivalence tests

In order to be more rigorous in declaring the null hypothesis of equality is accepted (in contrary to declaring the alternative cannot be rejected), equivalence tests in the form of Two-One-Sided t-Tests (TOST) were conducted (Lakens, 2017). Following the usual practice, the smallest effect size of interest is set at 30% to characterize the upper and lower equivalence bounds. All raw p-values from TOST were corrected for multiple comparisons errors by FDR corrections.

### Two-way ANOVA

Data from mouse models were analyzed by two-way ANOVA, followed by Sidak’s multiple comparison test. mRNA expression was considered significantly different from SAAR only if the fold-change was > 2 and p-values were < 0.05. P-values for interaction between the two independent variables was represented as μ_int,_ and pairwise comparisons were represented by μ.

### Clinical and epidemiological data

Biomarker data were log-transformed before analysis due to non-normal distribution. Two regression analyses were performed. First, we investigated the association of each of the plasma amino acids (Met, tCys, and Ser) with plasma triglycerides in crude models and models adjusted for age, sex, and BMI (log[triglycerides] ∼ log[amino acid] + age + sex + BMI). Second, we investigated whether plasma amino acids changed with the occurrence of MetS criteria in crude models and models adjusted for age and sex (log[amino acid] ∼ MetS criteria + age + sex). BMI was not considered for the second regression analysis as obesity is one of the five criteria for MetS. Since regression models on plasma amino acids and triglycerides were run on log-transformed values, estimates were expressed as a percentage increase in triglyceride concentration per percentage increase in the amino acids’ concentrations. Estimates from regression models between plasma amino acids and MetS criteria were expressed as a percentage change in amino acids with an increase in one MetS criterion. All data were expressed as geometric means (gM) and geometric standard deviations (gSD). In controlled-feeding studies, differences in plasma Ser between groups were analyzed using linear mixed model regression with Ser as the outcome and group, visit, and their interaction term (group × visit) as predictors. Models were adjusted for baseline differences.

## Acknowledgments

**SNN** received funding from the Orentreich Foundation for the Advancement of Science for the projects ASL18 and ASL32; **VK** and **JS** received institutional support from the project RVO-VFN 64165 of the Ministry of Health of the Czech Republic, and from the program of the Charles University Cooperatio-Metabolic Medicine/Endocrinology, Diabetology, and Metabolism; **GPA** received funding from the Orentreich Foundation for the Advancement of Science for the projects ASL21 and ASL 24; **KJV** received funding from the Research Council of Norway and the University of Oslo for the project ES528805. **VM** received funding from the National Institute of Environmental Health Science for the project P30ES023515.

## Contributions

**SNN** – Conceptualized the entire study, designed rat experiments, analyzed data from rat and mouse studies, interpreted data, wrote manuscript, initiated collaborations, and managed all aspects of the project; **MP** – Epidemiological study design, randomized selection of samples from the original cohort for the present study, initial modeling; **JS** and **VK** – Epidemiological study design, data acquisition, preliminary analyses of the parent epidemiological study; **Bø**- Designed diets for both controlled-feeding studies and involved in the design of Study 1; **CT** - Developed methods for quantitation of Ser, collected and analyzed data in Study 2. **NEB**- Analyzed Ser in Study 1 and collected data from both controlled feeding studies; **HR** - Design both pilot studies, co-developed methods for Ser quantification; involved in developing hypotheses for the controlled feeding studies; **KJV** – Designed, supervised, and collected data from the controlled feeding studies. **TO** – Collected and analyzed data from controlled feeding studies, designed and managed Study 1, analyzed data from the epidemiological study; **VM** and **JPR** – developed and analyzed rat data for equivalence tests and regression modeling for dose-response; **AKS** and **WM** – plasma amino acid analysis from rat experiments; **GPA** – Designed and lead mouse-model studies; **DC** – Animal husbandry and tissue harvesting from mouse-model experiments; **VLM** - Animal husbandry and tissue harvesting from rat experiments; **DALM** - Animal husbandry for rat studies, tissue harvesting, and all molecular and biochemical analysis of all specimens collected from rats and mice.

