## Supplementary Tables for "Cysteine Restriction-Specific Effects of Sulfur Amino Acid Restriction on Lipid Metabolism"

| **Supplementary Table 1a. Composition of Experimental Diets Used in Rat Experiments ^a^** | | |
| --- | --- | --- |
| **Ingredient** |  | **Composition ((gm% (Kcal%))** |
| Protein |  | 14 (14) |
| Carbohydrate |  | 70 (69) |
| Fat |  | 8 (18) |
| Kcal/gm^b^ |  | 4.1 |
| L-Arginine |  | 1.12 (1.11) |
| L-Histidine-HCl-H_2_O |  | 0.33 (0.33) |
| L-Isoleucine |  | 0.82 (0.81) |
| L-Leucine |  | 1.11 (1.10) |
| L-Lysine |  | 1.44 (1.42) |
| L-Phenylalanine |  | 1.16 (1.15) |
| L-Threonine |  | 0.82 (0.81) |
| L-Tryptophan |  | 0.18 (0.18) |
| L-Valine |  | 0.82 (0.81) |
| Glycine |  | 2.33 (2.30) |
| ***DL-Methionine+L-Cysteine+L-Glutamic Acid* ^c^** |  | ***0.35 (3.51)*** |
| Corn Starch |  | 36.11 (35.65) |
| Maltodextrin |  | 12.5 (12.34) |
| Sucrose |  | 20.0 (19.74) |
| Cellulose |  | 5.0 (0) |
| Corn Oil |  | 8.0 (17.77) |
| Mineral Mix S10001 |  | 3.5 (0) |
| Vitamin Mix V10001 |  | 1.00 (0.99) |
| Choline Bitrartrate |  | 2 (0) |
| **^a^** Nutrient composition in this table is common for all the diets used in rat cohorts  **^b^** Energy density was determined using Atwater factor system  **^c^** *See Table 1b for specific concentrations of Met, Cys, and Glu in individual diets* | | |

| **Supplementary Table 1b. Composition of Experimental Diets Used in Rat Experiments** | | | | | | | | |
| --- | --- | --- | --- | --- | --- | --- | --- | --- |
| **Diet** |  | **Research Diets (Catalog No.)** |  | **DL-Methionine (gm% (Kcal%))** |  | **L-Cysteine (gm% (Kcal%))** |  | **L-Glutamic Acid (gm% (Kcal%))** |
| CD |  | A15021902 |  | 0.86 (0.85) |  | 0 (0) |  | 2.7 (2.67) |
| SAAR |  | A15021901 |  | 0.17 (0.17) |  | 0 (0) |  | 3.39 (3.35) |
| MR1 |  | A15070801 |  | 0.17 (0.17) |  | 0.5 (0.49) |  | 2.89 (2.86) |
| MR2 |  | A15070802 |  | 0.10 (0.09) |  | 0.5 (0.49) |  | 2.96 (2.91) |
| MR3 |  | A15070803 |  | 0.07 (0.07) |  | 0.5 (0.49) |  | 2.99 (2.96) |
| MR4 |  | A15070804 |  | 0.05 (0.05) |  | 0.5 (0.49) |  | 3.01 (2.96) |
| CR1 |  | A15021903 |  | 0.07 (0.07) |  | 0.5 (0.49) |  | 2.99 (2.95) |
| CR2 |  | A15021904 |  | 0.07 (0.07) |  | 0.25 (0.24) |  | 3.24 (3.20) |
| CR3 |  | A15021905 |  | 0.07 (0.07) |  | 0.12 (0.12) |  | 3.37 (3.33) |
| CR4 |  | A15021906 |  | 0.07 (0.07) |  | 0.06 (0.06) |  | 3.43 (3.38) |
| CR5 |  | A15021907 |  | 0.07 (0.07) |  | 0.03 (0.03) |  | 3.46 (3.42) |

| **Supplementary Table 2. Composition of Experimental Diets Used in Mouse Experiments** | | | | | | | |
| --- | --- | --- | --- | --- | --- | --- | --- |
| **Ingredient** | **Composition in 10% Kcal**  **fat diets (gm%[kCal%])** | | |  | **Composition in 60% Kcal**  **fat diets (gm%[kCal%])** | | |
|  | **CD** |  | **SAAR** |  | **CD** |  | **SAAR** |
|  | **A14040402^a^** |  | **A14040401^a^** |  | **A14032002^a^** |  | **A14032001^a^** |
| Protein | 13 (14) |  | 13 (14) |  | 17 (13) |  | 17 (13) |
| Carbohydrate | 74 (76) |  | 74 (76) |  | 36 (28) |  | 36 (28) |
| Fat | 4 (10) |  | 4 (10) |  | 35 (60) |  | 35 (60) |
| Kcal/gm^b^ | 3.9 |  | 3.9 |  | 5.3 |  | 5.3 |
| L-Arginine | 1.09 (1.12) |  | 1.09 (1.12) |  | 1.48 (1.13) |  | 1.48 (1.13) |
| L-Histidine-HCl-H_2_O | 0.32 (0.32) |  | 0.32 (0.32) |  | 0.44 (0.33) |  | 0.44 (0.33) |
| L-Isoleucine | 0.8 (0.82) |  | 0.8 (0.82) |  | 1.09 (0.83) |  | 1.09 (0.83) |
| L-Leucine | 1.08 (1.1) |  | 1.08 (1.1) |  | 1.47 (1.1) |  | 1.47 (1.1) |
| L-Lysine | 1.4 (1.45) |  | 1.4 (1.45) |  | 1.91 (1.45) |  | 1.91 (1.45) |
| **DL-Methionine** | **0.86 (0.87)** |  | **0.12 (0.12)** |  | **0.86 (0.65)** |  | **0.12 (0.1)** |
| **L-Cysteine** | **0 (0)** |  | **0 (0)** |  | **0 (0)** |  | **0 (0)** |
| L-Phenylalanine | 1.13 (1.15) |  | 1.13 (1.15) |  | 1.53 (1.15) |  | 1.53 (1.15) |
| L-Threonine | 0.8 (0.82) |  | 0.8 (0.82) |  | 1.09 (0.83) |  | 1.09 (0.83) |
| L-Tryptophan | 0.17 (0.17) |  | 0.17 (0.17) |  | 0.24 (0.18) |  | 0.24 (0.18) |
| L-Valine | 0.8 (0.82) |  | 0.8 (0.82) |  | 1.09 (0.83) |  | 1.09 (0.83) |
| **L-Glutamic Acid** | **2.7 (2.77)** |  | **3.44 (3.55)** |  | **2.7 (2.05)** |  | **3.43 (2.6)** |
| Glycine | 2.26 (2.32) |  | 2.26 (2.32) |  | 3.08 (2.33) |  | 3.08 (2.33) |
| **Corn Starch** | **41.17 (42.41)** |  | **41.17 (42.41)** |  | **0 (0)** |  | **0 (0)** |
| **Maltodextrin** | **12.12 (12.49)** |  | **12.12 (12.49)** |  | **8.68 (6.55)** |  | **8.68 (6.55)** |
| Dextrose | 4.85 (5) |  | 4.85 (5) |  | 6.62 (5) |  | 6.62 (5) |
| Sucrose | 14.55 (14.99) |  | 14.55 (14.99) |  | 19.85 (15) |  | 19.85 (15) |
| Cellulose | 4.85 (0) |  | 4.85 (0) |  | 6.62 (0) |  | 6.62 (0) |
| **Lard** | **0 (0)** |  | **0 (0)** |  | **28.98 (49.28)** |  | **28.98 (49.28)** |
| Corn Oil | 4.46 (10.34) |  | 4.46 (10.34) |  | 6.09 (10.35) |  | 6.09 (10.35) |
| Mineral Mix S10001 | 3.39 (0) |  | 3.39 (0) |  | 4.63 (0) |  | 4.63 (0) |
| Vitamin Mix V10001 | 0.97 (1) |  | 0.97 (1) |  | 1.32 (1) |  | 1.32 (1) |
| Choline Bitrartrate | 0.19 (0) |  | 0.19 (0) |  | 0.26 (0) |  | 0.26 (0) |
| ^a^ Research Diets catalog number  ^b^ Energy density was determined using Atwater factor system | | | | | | | |

| **Supplementary Table 3. Effect of Gender on the Molecular Mechanisms of SAAR-induced Changes in Lipid Metabolism** | | | | | | | |
| --- | --- | --- | --- | --- | --- | --- | --- |
|  | 10% Kcal Fat Diets | | |  | 60% Kcal Fat Diets | | |
|  | Fold Change in Males (SAAR/CD) | Fold Change in Females (SAAR/CD) | Ratio of Fold Change (Males/Females) |  | Fold Change in Males (SAAR/CD) | Fold Change in Females (SAAR/CD) | Ratio of Fold Change ( Males/Females ) |
| Decrease in Glutathione | 0.35 | 0.42 | 0.85**^n.s.^** |  | 0.17 | 0.26 | ***0.67****** |
| Increase in NRF2 | 3.65 | 3.97 | 0.92**^n.s.^** |  | 5.97 | 3.69 | ***1.62**** |
| Increase in PHGDH | 5.46 | 11.45 | 0.48**^n.s.^** |  | 11.56 | 5.06 | ***2.28**** |
| Increase in PCK2 | 24.55 | 23.68 | 1.04**^n.s.^** |  | 100.15 | 96.54 | 1.04**^n.s.^** |
| Decrease in Triglycerides | 0.87 | 0.89 | 0.97**^n.s.^** |  | 0.69 | 0.71 | 0.98**^n.s.^** |
| Decrease in Perigonadal adipose depot weight | 0.67 | 1.00 | ***0.67****** |  | 0.44 | 0.96 | ***0.46****** |
| Note: Experiments 3, 4, and 5 (for details see section on methods) were conducted at different time points but under the same animal husbandry conditions, with similar diets, and in the same animal facility. For data analysis purposes all three experiments were considered as one. Fold-changes and asterisks in columns 3 and 6 represent μ_int_-values obtained from two-way ANOVA, considering diet and gender as independent variables. | | | | | | | |

| **Supplementary Table 4. Effect of Age-at-onset on the Molecular Mechanisms of SAAR-induced Changes in Lipid Metabolism** | | | | | | | |
| --- | --- | --- | --- | --- | --- | --- | --- |
|  | Males | | |  | Females | | |
|  | Fold Change in Young (SAAR/CD) | Fold Change in Adult  (SAAR/CD) | Ratio of Fold Change (Y/A) |  | Fold Change in Young (SAAR/CD) | Fold Change in Adult (SAAR/CD) | Ratio of Fold Change (Y/A) |
| Decrease in Glutathione | 0.35 | 0.43 | 0.82**^n.s.^** |  | 0.42 | 0.71 | **0.59***** |
| Increase in NRF2 | 3.65 | 2.61 | 1.40**^n.s.^** |  | 3.97 | 1.36 | **2.92****** |
| Increase in PHGDH | 5.46 | 4.34 | 1.26**^n.s.^** |  | 11.45 | 4.87 | 2.35**^n.s.^** |
| Increase in PEPCK-M | 24.55 | 55.00 | 0.45**^n.s.^** |  | 23.68 | 7.75 | **3.05******* |
| Decrease in Triglycerides | 0.87 | 0.87 | 1.00**^n.s.^** |  | 0.89 | 0.99 | 0.90**^n.s.^** |
| Decrease in Perigonadal adipose depot weight | 0.67 | 0.68 | 1.00**^n.s.^** |  | 1.00 | 0.64 | 1.58**^n.s.^** |
| Note: Experiments 3, 4, and 5 (for details, see the section on methods) were conducted at different time points but under the same animal husbandry conditions, with similar diets, and in the same animal facility. For data analysis purposes, all three experiments were considered as one. Fold-changes and asterisks in columns 3 and 6 represent μ_int_-values obtained from two-way ANOVA, considering diet and age-at-onset as independent variables. | | | | | | | |

| **Supplementary Table 5: Epidemiological Study Population Characteristics** | | | | | | | |
| --- | --- | --- | --- | --- | --- | --- | --- |
| **Characteristic** | **0 MetS criteria** | **1 MetS criteria** | **2 MetS criteria** | **3 MetS criteria** | **4 MetS criteria** | **5 MetS criteria** | **P_trend_** |
| Sample size (n) | 50 | 50 | 50 | 50 | 50 | 57 | - |
| Age (y) | 38.5 (1.42) | 47.1 (1.34) | 51.3 (1.17) | 52.9 (1.16) | 53.4 (1.16) | 54.1 (1.11) | < 0.001 |
| Males, n (%) | 16 (32) | 23 (46) | 30 (60) | 37 (74) | 38 (76) | 41 (71.9) | < 0.001 |
| BMI (kg/m^2^) | 22 (1.09) | 24.6 (1.14) | 26.3 (1.11) | 28.3 (1.14) | 29.2 (1.15) | 30.6 (1.14) | < 0.001 |
| Triacylglycerols  (mmol/l) | 0.89 (1.38) | 1.01 (1.45) | 1.18 (1.45) | 1.83 (1.43) | 2.34 (1.39) | 3.17 (1.54) | < 0.001 |
| CAD patient, n (%) | 2 (4) | 8 (16) | 15 (30) | 24 (48) | 28 (56) | 41 (71.9) | < 0.001 |
| Smoker, n (%) | 13 (26) | 12 (24) | 6 (12) | 13 (26) | 5 (10) | 6 (10.5) | < 0.001 |
| Former smoker, n (%) | 8 (16) | 10 (20) | 17 (34) | 21 (42) | 31 (62) | 41 (71.9) | < 0.001 |
| Note: Continuous variables are presented as geometric mean (geometric standard deviation). Categorical variables are presented as n (%). | | | | | | | |

| **Supplementary Table 6: Associations of Plasma Amino Acids with Triglycerides and MetS Criteria** | | | | | |
| --- | --- | --- | --- | --- | --- |
| a) Plasma amino acids and triglycerides | | | | | |
|  | n | Unadjusted model | | Adjusted model^a^ | |
| Amino Acids | | Estimate^b^ (95 % CI) | p-value | Estimate^b^ (95 % CI) | p-value |
| Met | 307 | -0.001 (-0.281,0.279) | 0.994 | -0.066 (-0.3,0.168) | 0.578 |
| tCys | 307 | 1.338 (0.997,1.779) | < 0.001 | 0.546 (0.125,0.968) | 0.011 |
| Ser | 287 | -0.397 (-0.619,-0.174) | 0.001 | -0.302 (-0.496,-0.108) | 0.002 |
| tCys/Ser | 287 | 0.57 (0.37, 0.769) | < 0.001 | 0.337 (0.155, 0.52) | < 0.001 |
| b) Plasma amino acids and MetS criteria | | | | | |
|  | | Unadjusted model | | Adjusted model^c^ | |
| Amino Acids | | β-estimate^d^ (confidence intervals) | p-value | β-estimate^d^ (confidence intervals) | p-value |
| Met | 307 | 0.939 (-0.615,2.518) | 0.237 | 0.881 (-0.883,2.677) | 0.329 |
| tCys | 307 | 4.104 (3.143,5.075) | < 0.001 | 2.154 (1.18,3.137) | < 0.001 |
| Ser | 287 | -2.145 (-4.123,-0.127) | 0.037 | -2.563 (-4.729,-0.348) | 0.024 |
| tCys/Ser | 287 | 5.306 (3.08,7.579) | < 0.001 | 4.294 (1.862,6.785) | < 0.001 |
| ^a^ Adjusted for age, gender, and BMI  ^b^ β-estimates represent % change in triglycerides per 1 % change in the amino acid concentration.  ^c^ Adjusted for age and gender  ^d^ β-estimates represent % change in the amino acid concentration per increase in each MetS criteria | | | | | |

| **Supplementary Table 7. Dietary Sulfur Amino Acid and Fat Composition in Short-term Feeding Studies** | | | | | | |
| --- | --- | --- | --- | --- | --- | --- |
|  | SAA intake ^a^ (g/day(mg/Kg BW/day)) | | Fat intake (% total energy) | | Sample size | |
|  | Men | Women | PUFA | SFA | Males | Females |
| Study 1 | | | | | | |
| SAA_low_ | -- | 1.5 (17.7) | 7.2 | 3.9 | 0 | 7 |
| SAA_high_ | -- | 5.5 (71.5) | 7.2 | 3.9 | 0 | 6 |
| Study 2 | | | | | | |
| SAA_low+PUFA_ | 1.19 (15.3) | 0.93 (14.2) | 10 | 5.5 | 2 | 5 |
| SAA_high+SFA_ | 6.01 (82.0) | 5.75 (87.9) | 3.3 | 13.5 | 2 | 5 |
| ^a^ represents the intake of Met and Cys together | | | | | | |

| **Supplementary Table 8. Changes in Plasma Ser in Short-term Controlled Feeding Studies** | | | | |
| --- | --- | --- | --- | --- |
| Diet | Plasma Ser (mmol/L, Mean±SEM) | | | p-value for time-related (day 0-7) changes between the diet groups |
|  | Day 0 | Day 3 | Day 7 |  |
| SAA_low_ | 100.85±9.23 | 102.79±6.84 | 96.89±7.72 | 0.94 |
| SAA_high_ | 103.70±6.81 | 107.07±6.64 | 100.72±9.97 |  |
| SAA_low+PUFA_ | 105.82±8.06 | 105.24±6.42 | 118.02±8.49 | 0.018 |
| SAA_high+SFA_ | 101.21±8.55 | 91.96±7.47 | 96.12±6.59 |  |

| **Supplementary Table 9. Confirmation of Sulfur Amino Acid Concentration in Experimental Diets Used for Rat Cohorts** | | | | |
| --- | --- | --- | --- | --- |
| Diet |  | Methionine (Detected/Expected) |  | Cysteine (Detected/Expected) |
| CD |  | **0.783**/0.86 |  | **<0.01**/0 |
| SAAR |  | **0.162**/0.17 |  | **<0.01**/0 |
| MR1 |  | **0.164**/0.17 |  | **0.33**/0.5 |
| MR2 |  | **0.109**/0.1 |  | **0.36**/0.5 |
| MR3 |  | **0.071**/0.07 |  | **0.337**/0.5 |
| MR4 |  | **0.056**/0.05 |  | **0.348**/0.5 |
| CR1 |  | **0.075**/0.07 |  | **0.338**/0.5 |
| CR2 |  | **0.073**/0.07 |  | **0.178**/0.25 |
| CR3 |  | **0.079**/0.07 |  | **0.101**/0.125 |
| CR4 |  | **0.075**/0.07 |  | **0.051**/0.062 |
| CR5 |  | **0.073**/0.07 |  | **0.033**/0.031 |
| Note: Dietary analysis was performed at Covance | | | | |

| **Supplementary Table 10. Experimental Details of Protein and mRNA Quantification** | | | | | | | | | |
| --- | --- | --- | --- | --- | --- | --- | --- | --- | --- |
| a) Enzyme-linked immunosorbent assays | | | | | | | | | |
| Plasma marker | | Vendor | | | Catalog# | | Comments | | |
| Igf-1 | | R&D Systems | | | MG100 | | Mouse/Rat Kit | | |
| Adiponectin | | R&D Systems | | | RRP300 | | Rat Kit | | |
| Fgf21 | | R&D Systems | | | RRP300 | | Rat Kit | | |
| Leptin | | R&D Systems | | | MOB00 | | Mouse/Rat Kit | | |
| b) Western blot details | | | | | | | | | |
| *Protein of interest* | *Antibody source* | | | | *Dilution* | | | *Incubation conditions* | |
|  | *Primary* | | | *Secondary* | *Primary* | *Secondary* | | *Primary* | *Secondary* |
| Phgdh (Rat) | Cell Signaling  13428 | | | Cell Signaling  7074 | 1:1000 | 1:7500 | | 5% milk O/N @ 4^o^C | 5% milk 1 hr @ RT |
| Phgdh (Mouse) | Cell Signaling  13428 | | | Cell Signaling  7074 | 1:1000 | 1:7500 | | 5% milk O/N @ 4^o^C | 5% milk 1 hr @ RT |
| Pepck-M (Rat) | Cell Signaling  6924 | | | Cell Signaling  7074 | 1:1000 | 1:7500 | | 5% milk O/N @ 4^o^C | 5% milk 1 hr @ RT |
| Pepck-M (Mouse) | Cell Signaling  6924 | | | Cell Signaling  7074 | 1:1000 | 1:7500 | | 5% milk O/N @ 4^o^C | 5% milk 1 hr @ RT |
| Nrf2 (Mouse) | Cell Signaling  20733 | | | Cell Signaling  7074 | 1:1000 | 1:7500 | | 5% milk O/N @ 4^o^C | 5% milk 1 hr @ RT |
| β-Actin | MilliporeSigma  A5441 | | | Bio-Rad  1706516 | 1:20000 | 1:20000 | | 5% milk 30 min @ RT | 5% milk 30 min @ RT |
| Vinculin | Proteintech  66305-1-lg | | | Bio-Rad  1706516 | 1:10000 | 1:20000 | | 5% milk 30 min @ RT | 5% milk 30 min @ RT |
| c) Assay details for mRNA quantification | | | | | | | | | |
| *Gene of interest* | | | *TaqMan assay ID* | | | | | | |
| *Phgdh* | | | Rn01534200_g1 | | | | | | |
| *Pck1* | | | Rn01529014_m1 | | | | | | |
| *Pck2* | | | Rn03648110_m1 | | | | | | |
| *G6pc* | | | Rn00689876_m1 | | | | | | |
| *β2M* | | | Rn00560865_m1 | | | | | | |

**Figure Legends**

**Figure 1: Methionine restriction and cysteine restriction exert discrete effects on morphometry and plasma hormone concentrations.** Eight-week-old male F344 rats were fed CD, SAAR, MR (MR1, MR2, MR3, and MR4), and CR (CR1, CR2, CR3, CR4, and CR5) diets for 12 weeks. Changes in growth rate (a), food intake (b), plasma Igf1 (c), plasma Fgf21 (d), and plasma leptin (e), were dependent on MR, while changes in plasma adiponectin (f) were dependent on CR. *Note*: n = 16 for CD and SAAR groups, n=8/group for all other groups; μ indicates the mean differences between CD and SAAR groups. MR_β_ and CR_β_ indicate whether the dose-response (regression coefficients) occurred or not with MR and CR, respectively. Bars and error bars represent the means and standard error of means. Asterisks indicate the significance of statistical differences, i.e., p<0.05 (*), p<0.01(**), p<0.001(***), p<0.0001 (****), and n.s. (not significant).

**Figure 2: Methionine restriction and cysteine restriction induce distinct changes in plasma amino acid concentrations.** Although SAAR changed plasma concentrations of glutamic acid (a), glycine (b), and lysine (c) concentrations, these amino acids did not show dose-response to either MR or CR. A strong dose-response was exhibited by plasma histidine (d), methionine (e), phenylalanine (f), serine (g), threonine (h), and tryptophan (i) to CR. *Note*: n = 16 for CD and SAAR groups, n=8/group for all other groups; μ indicates the mean differences between CD and SAAR groups.

**Figure 3: Cysteine restriction, but not methionine restriction, induces hepatic *de novo* Ser biosynthesis.** CR but not MR increases hepatic serine concentrations (a), *Phgdh* mRNA (b and c), and Phgdh protein levels (d and e). *Note*: in panel a, n=16 for CD and SAAR groups, n=8/group for all other groups; in all other panels, n = 4-8/group; in panel d, only the diets with the highest and lowest concentration of Met (MR1 and MR4) and Cys (CR1 and CR5) were probed.

**Figure 4: Cysteine restriction increases serinogenesis at the expense of glyceroneogenesis.** MR but not CR decreased blood glucose (a). Neither MR nor CR altered the mRNA expression of *G6pc* (b and c) and *Pck1* (d and e). CR but not MR increased the hepatic mRNA expression of *Pck2* (f and g) and its protein Pepck-M (h and i), and decreased hepatic glycerol-3-phosphate concentration (j). *Note*: in panel a, n = 14 for CD and SAAR groups, n=8/group for all other groups; in all other panels, n = 4-8/group;

**Figure 5: SAAR-induced changes in molecular markers of serinogenesis and lipid metabolism are dependent on gender, dietary fat content, and age-at-onset.** Young (eight-week-old, solid and empty bars) and adult (eighteen-month-old, hatched bars) male (blue bars) and female (red bars) C57BL/6J mice were fed CD (empty bars) and SAAR (solid bars) diets with either 10% fat or 60% fat for at least three months. Statistical significance of SAAR-induced changes in hepatic glutathione (a-c), hepatic protein expressions of Nrf2 (d-f), Phgdh (g-i), and Pck2 (j-l), plasma triglycerides (m-o), and perigonadal adipose tissue weights (p-r) are indicated by μ. The dependence of SAAR-induced changes in these markers on dietary fat and gender are indicated by μ_int_. Note: n=8/group; PGAT-perigonadal adipose tissue; bars and error bars indicate means and standard error of means. Asterisks indicate the significance of statistical differences, i.e., p<0.05 (*), p<0.01(**), p<0.001(***), p<0.0001 (****), and n.s. (not significant).

**Figure 6: Plasma tCys, Ser, and tCys/Ser correlate with plasma triglycerides and the number of MetS criteria in humans**. Log-transformed plasma concentrations of Met (a and e), tCys (b and f), Ser (c and g), and tCys/Ser (d and h) were plotted against log-transformed triglycerides (a-d) and MetS criteria (e-h), respectively. Red and blue lines in figures a-d represent unadjusted and adjusted (for age, gender, and body mass index) regression lines, respectively. Note: n = 307 for plasma Met and tCys and n=287 for plasma Ser; box plots represent the distribution of unadjusted amino acid concentrations within each category of MetS. Numbers in parenthesis after β_un_ and β_ad_ represent regression coefficients from unadjusted and adjusted models, respectively; asterisks represent the significance of statistical differences, i.e., p<0.05 (*), p<0.01(**), p<0.001(***), p<0.0001 (****), and n.s. (not significant).

**Figure 7: Plasma Ser levels are amenable to low SAA diets depending on dietary fatty acid composition** a) Change in plasma serine in overweight/obese women (n = 13) on a 7-day pilot dietary intervention trial with low (1.5 g/day, SAA_low_) or high (5.5 g/day, SAA_high_) sulfur amino acid content. P-values represent group x time interactions (change in plasma serine over time between groups) and were computed with linear mixed models adjusted for baseline levels of serine and correlated observations among subjects. b) Change in plasma serine in normal-weight individuals (n = 14) participating in a 7-day pilot dietary intervention trial with high polyunsaturated fatty acid contents (10.6 E %/day for women and 10.9 E %/day for men) and low sulfur amino acid concentrations (0.93 g/day for women, 1.18 g/day for men, SAA_low+PUFA_) vs. a diet high in saturated fatty acids (13.5 E %/day for women, 13.3 E %/day for men) and high sulfur amino acids (5.75 g/day for women, 6.01 g/day for men, SAA_high+SFA_). Note: Black and blue dots represent mean Ser concentrations on high and low sulfur amino acid diets, respectively. Error bars represent confidence intervals.

**Supplementary Figures 1 and 2:** Two one-sides t-tests (TOST) were conducted to find which of the Met restricted (MR1, MR2, MR3, and MR4) and Cys restricted (CR1, CR2, CR3, CR4, and CR5) diets resulted in an equivalent response as the SAAR diet, in each of the six phenotypes tested (body weight, food intake, Igf1, Fgf21, leptin, and adiponectin). The X-axis represents the mean difference of each diet (CD [■, □], MR [●] and CR [○] diets) from the SAAR diet, horizontal lines on either side of the symbols represent 95% confidence intervals, vertical lines represent the 30% confidence bounds, and the asterisks represent the p-values obtained from TOST. Note: Although we did not get a significant p-value for growth rate, food intake, and Fgf21, we considered MR3 as the equivalent dose because it falls within the 30% confidence bounds (vertical lines) for most of the phenotypes.

**Summary Figure:** The proposed mechanism of CR-specific effects of SAAR on adipose metabolism. 1) Due to the lack of Cys in SAAR diets, it imposes both MR and CR. CR specifically results in decreased biosynthesis of the tripeptide glutathione 2) Lower hepatic glutathione increases the abundance of the transcription factor Nrf2, which translocates to the nucleus and induces the transcription of *Phgdh* 3) Increase in the transcription and translation of *Phgdh* results in higher hepatic serinogenesis (Ser biosynthesis from substrates other than glucose, i.e., from oxaloacetic acid) 4) Higher serinogenesis competes with glyceroneogenesis as both pathways use the same set of substrates, which results in lower fatty acid re-esterification. *Note*: broken arrows represent the presence of other biochemical intermediates not shown in the figure. Up and down arrows adjacent to the metabolic intermediates represent an increase and decrease, respectively. 3PG – 3-phosphoglycerate, OAA – Oxaloacetic acid, DHAP – Dihydroxyacetone phosphate, PEP-Phosphoenolpyruvate, FA – fatty acids.
