## Supplementary figures and images for "Cysteine Restriction-Specific Effects of Sulfur Amino Acid Restriction on Lipid Metabolism"

### Supplementary Figure 1

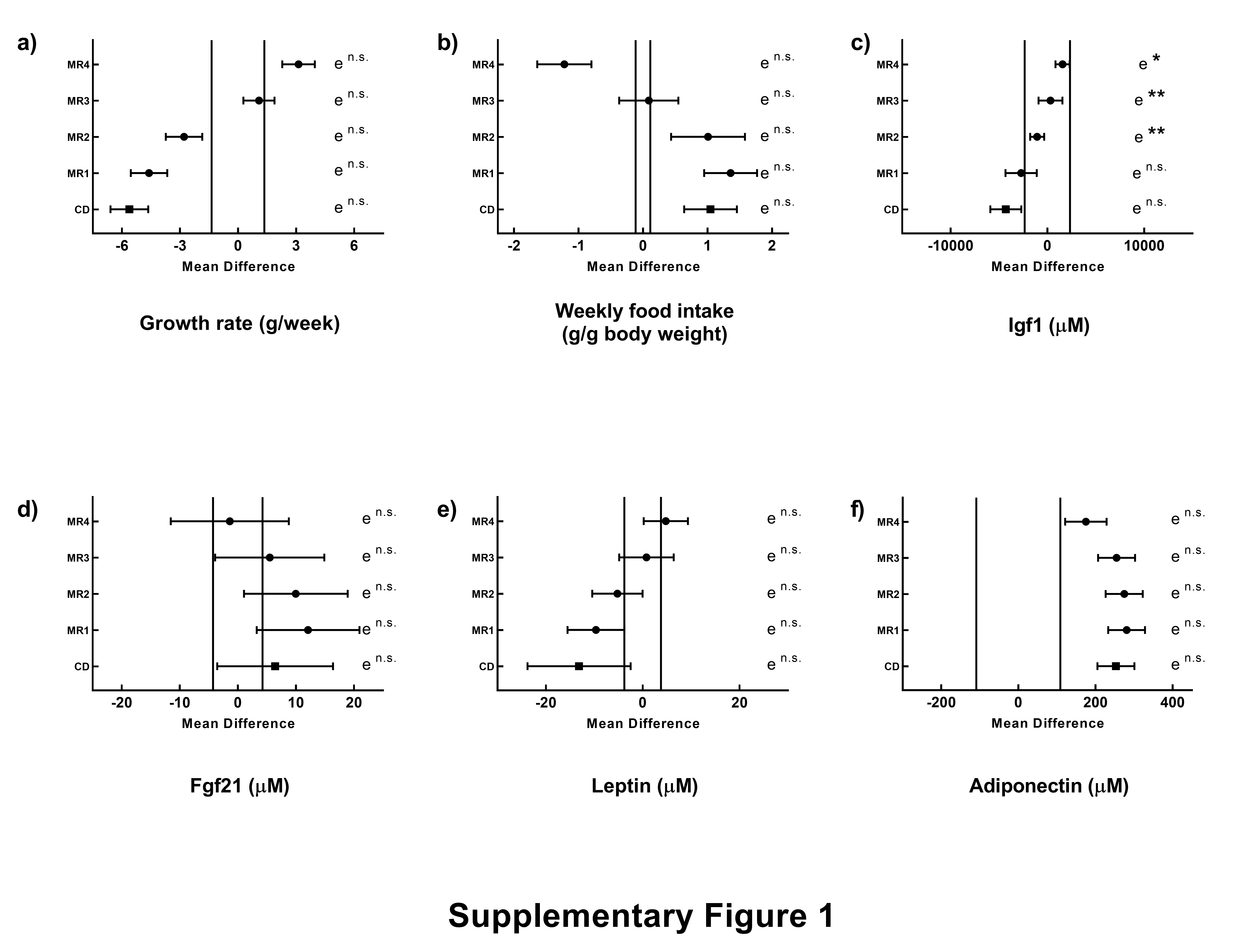

### Supplementary Figure 2

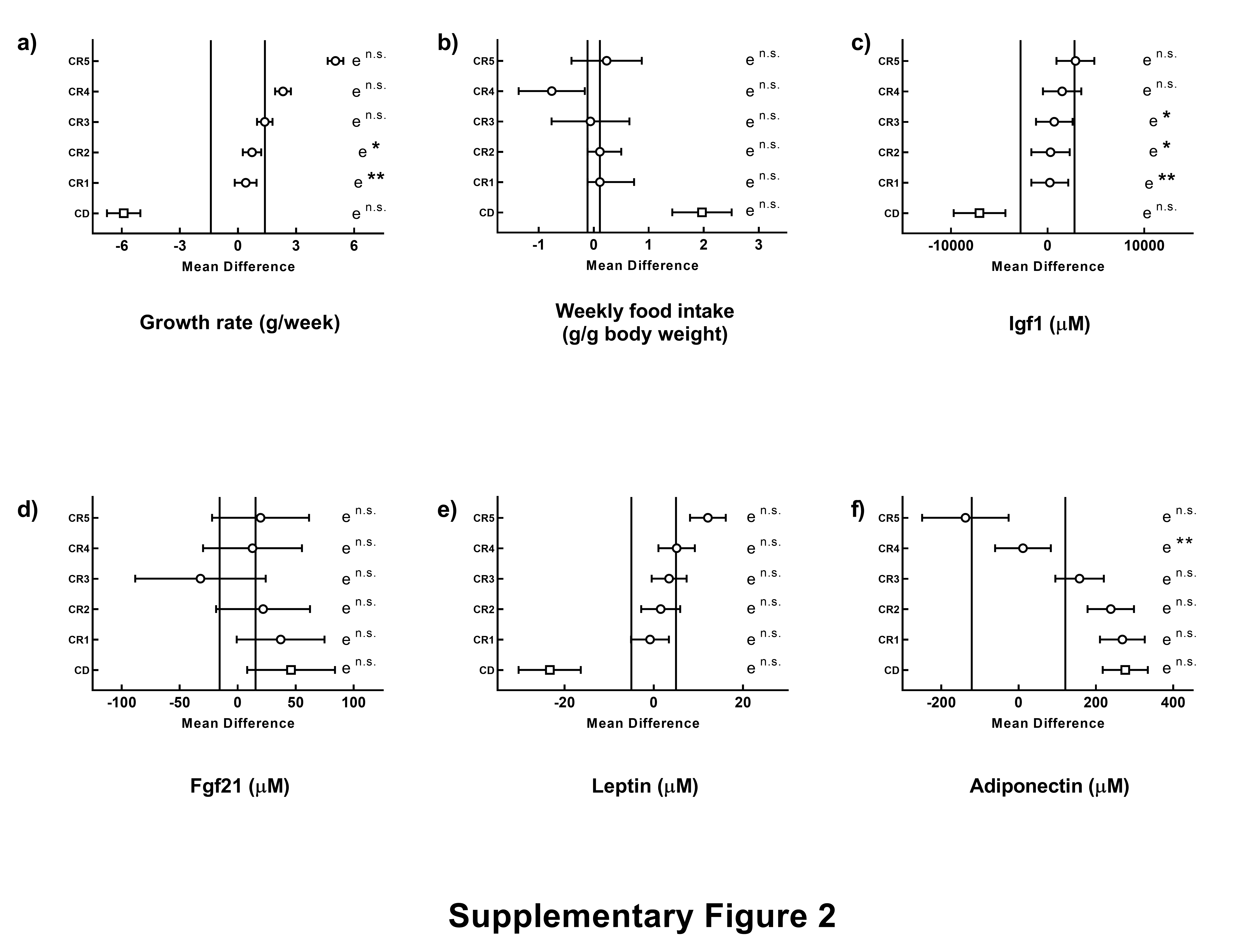
